# Cas9-generated auxotrophs of *Phaeodactylum tricornutum* are characterized by small and large deletions that can be complemented by plasmid-based genes

**DOI:** 10.1101/2020.04.12.038471

**Authors:** Samuel S. Slattery, Helen Wang, Daniel J. Giguere, Csanad Kocsis, Bradley L. Urquhart, Bogumil J. Karas, David R. Edgell

## Abstract

The model diatom *Phaeodactylum tricornutum* is an attractive candidate for synthetic biology applications. Development of auxotrophic strains of *P. tricornutum* would provide alternative selective markers to commonly used antibiotic resistance genes. Here, using CRISPR/Cas9, we show successful editing of genes in the uracil, histidine, and tryptophan biosynthetic pathways. Editing events are characterized by loss of heterozygosity and by the occurrence of large deletions of up to ~2.7-kb centred on the editing site. The uracil and histidine-requiring phenotypes can be complemented by plasmid-based copies of the intact genes after curing of the Cas9-editing plasmid. Growth of uracil auxotrophs on media supplemented with 5-FOA and uracil results in loss of the complementing plasmid, providing a facile method for plasmid curing with potential applications in strain engineering and CRISPR editing. Metabolomic characterization of uracil auxotrophs revealed changes in cellular orotate concentrations consistent with partial or complete loss of orotate phosphoribosyl transferase activity in knockout strains. Our results expand the range of *P. tricornutum* auxotrophic strains and demonstrate that auxotrophic complementation markers provide a viable alternative to traditionally used antibiotic selection markers. Plasmid-based auxotrophic markers should expand the range of genome engineering applications and provide a means for biocontainment of engineered *P. tricornutum* strains.

## Introduction

Photoautotrophic microalgae and cyanobacteria are emerging as alternative platforms for synthetic biology applications^1, 2^. One microalgae species of interest is the marine diatom *Phaeodactylum tricornutum*. A variety of plasmid-based genetic tools have been developed for *P. tricornutum* that facilitate basic molecular manipulations and expression of complex synthetic pathways^3–6^. We, and others, have developed plasmid-based and DNA-free CRISPR (clustered regularly interspaced palindromic repeats) reagents for targeted editing of the *P. tricornutum* chromosome using the Cas9 protein (CRISPR-associated protein 9)^6–10^. These plasmid-based tools and synthetic pathways are currently maintained by available antibiotic-based selections, including Zeocin™, phleomycin, nourseothricin, and blasticidin-S and their respective resistance genes, *Sh ble, nat*, and *bsr*^11–14^. Antibiotic-based selections can be prohibitively expensive for maintaining large-scale cultures and are problematic for applications such as the biosynthesis of products intended for human consumption^15–17^.

A viable alternative to antibiotics is the use of auxotrophic selective markers which require a strain engineered to have a loss of function mutation in a key enzyme of an essential biosynthetic pathway. Examples of commonly used auxotrophic strains in industrial and academic labs include uracil, histidine, and tryptophan auxotrophs^18–20^. Two approaches have been taken to generate *P. tricornutum* auxotrophs. First, uracil-requiring mutants were generated by random mutagenesis that resulted in the identification of the bi-functional uridine monophosphate synthase (PtUMPS) gene predicted to catalyze the conversion of orotate into uridine monophosphate (UMP)^21^. Biolistic transformation and chromosomal integration of the PtUMPs gene rescued the uracil-requiring phenotype. Second, Cas9 was used to knockout the PtUMPS gene to create uracil auxotrophs and the PtAPT gene encoding a predicted adenine phosphoribosyl transferase to create adenine auxotrophs^7^. However, direct selection of these auxotrophs via transformation with the corresponding complementation marker has not been explored and the generation of additional auxotrophic strains would facilitate development of new plasmid-based complementation markers.

Here, we used a plasmid-based editing strategy to generate knockouts in the uracil, histidine, and tryptophan biosynthesis pathways of *P. tricornutum* and show for the first time that plasmid-based copies of the intact PtUMPS and PtPRA-PH/CH genes can complement the uracil- and histidine-requiring phenotypes, respectively. Individual auxotroph strains are characterized by loss of heterozygosity at the edited alleles, and Nanopore sequencing of the edited population reveals large, heterogeneous deletions up to ∼ 2.7 kb. The uracil and histidine auxotrophs and their respective complementation markers are a potential alternative to antibiotic-based selection of plasmids in *P. tricornutum*. Our results also suggest a simple methodology to cure plasmids from uracil auxotrophs to enable strain and genome engineering.

## Results and Discussion

### Identification of Cas9 targets in biosynthetic pathway genes

We examined the KEGG predictions^22, 23^ based on the genome sequence of *P. tricornutum* to identify genes in the uracil and histidine biosynthetic pathways for Cas9 editing. We focused on these two pathways as uracil and histidine auxotrophy, and counter-selection strategies are commonly used in other model organisms. This approach identified the previously described bi-functional PtUMPS gene that is predicted to catalyze two steps in the uracil pathway — conversion of orotate to orotidine monophosphate (OMP), and conversion of OMP to uridine monophosphate (UMP) (Figure 1A)^21^. Proteins that are orthologs of characterized enzymes involved in histidine biosynthesis were also identified (Figure 2A). The PHATR_3140 gene, hereafter called PtPRA-PH/CH, encodes a predicted bifunctional protein that shares sequence similarity with the bacterial protein HisIE, and its plant counterpart HISN2^24, 25^. These proteins possess two functional domains that are homologous to the phosphoribosyl-ATP pyrophosphohydrolase (PRA-PH) and phosphoribosyl-AMP cyclohydrolase (PRA-CH) enzymes, respectively. PRA-PH and PRA-CH, alone or as a bifunctional protein, are predicted to catalyze two successive steps that occur early in the histidine biosynthesis pathway (Figure 2A). The PtIGPS gene encoding imidazole glycerol phosphate synthase (a HIS3 homolog) was found to be a duplicated gene in the *P. tricornutum* genome assembly and thus not prioritized as a Cas9 target.

**Figure 1.**
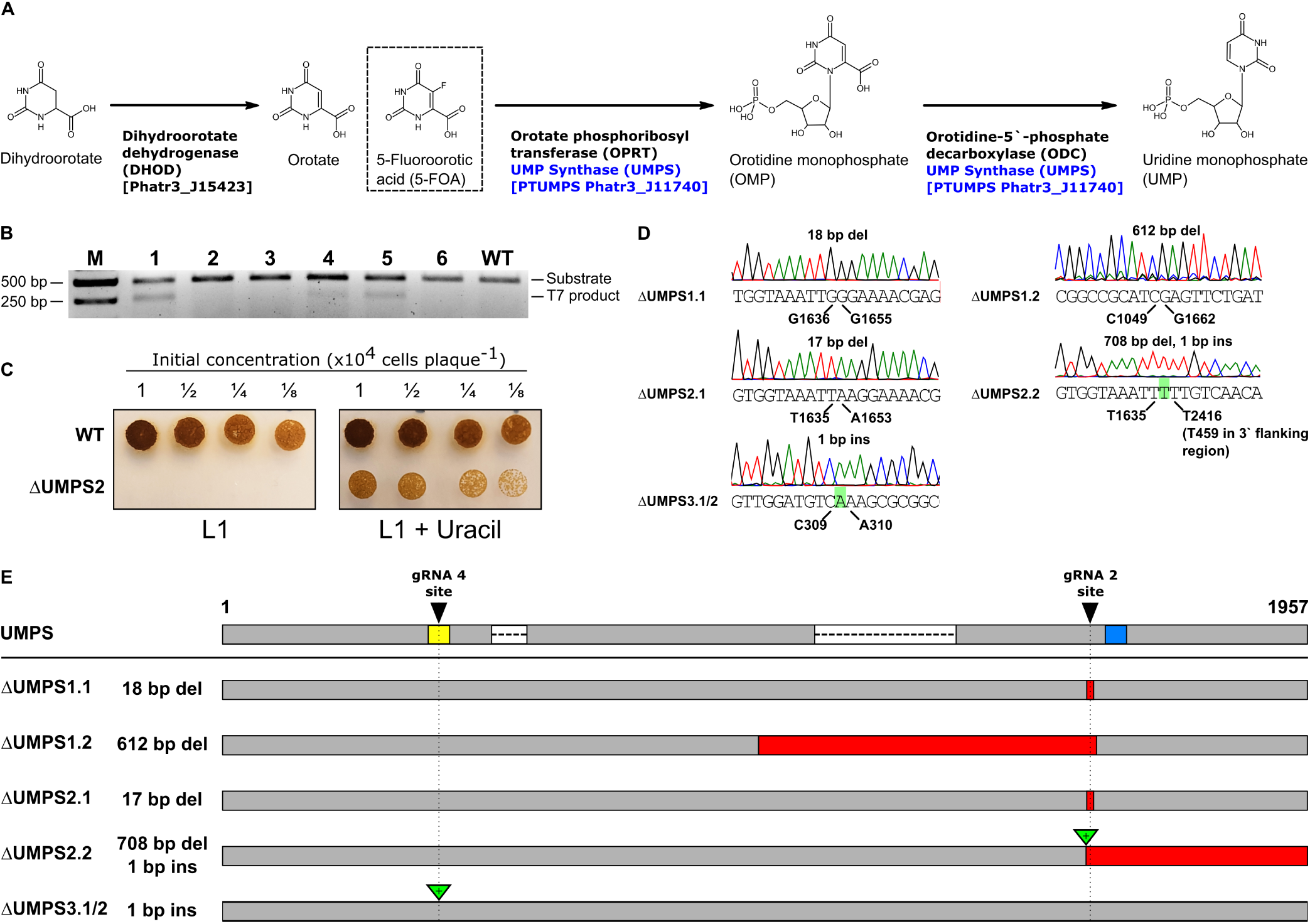
CRISPR-generated knockouts in the predicted *P. tricornutum* uracil biosynthesis pathway. (A) The predicted *P. tricornutum* biosynthesis pathway for conversion of carbonic acid to uracil and uridine triphosphate, with the PtUMPS enzyme highlighted in blue. The competitive inhibitor, 5-fluoroorotic acid (5-FOA), is also shown in a dashed box at the position where it enters the pathway. Abbreviated names for molecules and enzymes are indicated in parentheses, and the predicted corresponding *P. tricornutum* gene names are indicated in square brackets. (B) Example image of T7EI editing assay to screen exconjugants for potential editing events in the PtUMPS gene. Substrate indicates PtUMPS gene fragments amplified by the PCR, while T7 product indicates exconjugants with evidence of Cas9 editing. WT, wild-type *P. tricornutum* genomic DNA used in the T7EI editing assay. (C) Example of phenotypic screening of one PtUMPS knockout strain (ΔUMPS2) plated on L1 alone or L1 supplemented with uracil at the indicated dilution of initial concentration. (D) Sanger sequencing traces of characterized PtUMPS knockouts with the position (below trace) and type of insertion or deletion (above trace) indicated for each allele of the three strains. (E) Graphical map of the position and extent of indels for each of the three PtUMPS knockouts relative the wild-type UMPS gene (shown at top). Red rectangles indicate nucleotide deletions, green rectangles indicate nucleotide insertions, the yellow and blue rectangles on the WT gene indicate the position of the PtUMPS active sites (orotate phosphoribosyl transferase and orotidine-5’-phosphate decarboxylase), and the white rectangles with dashed lines represent introns.

**Figure 2.**
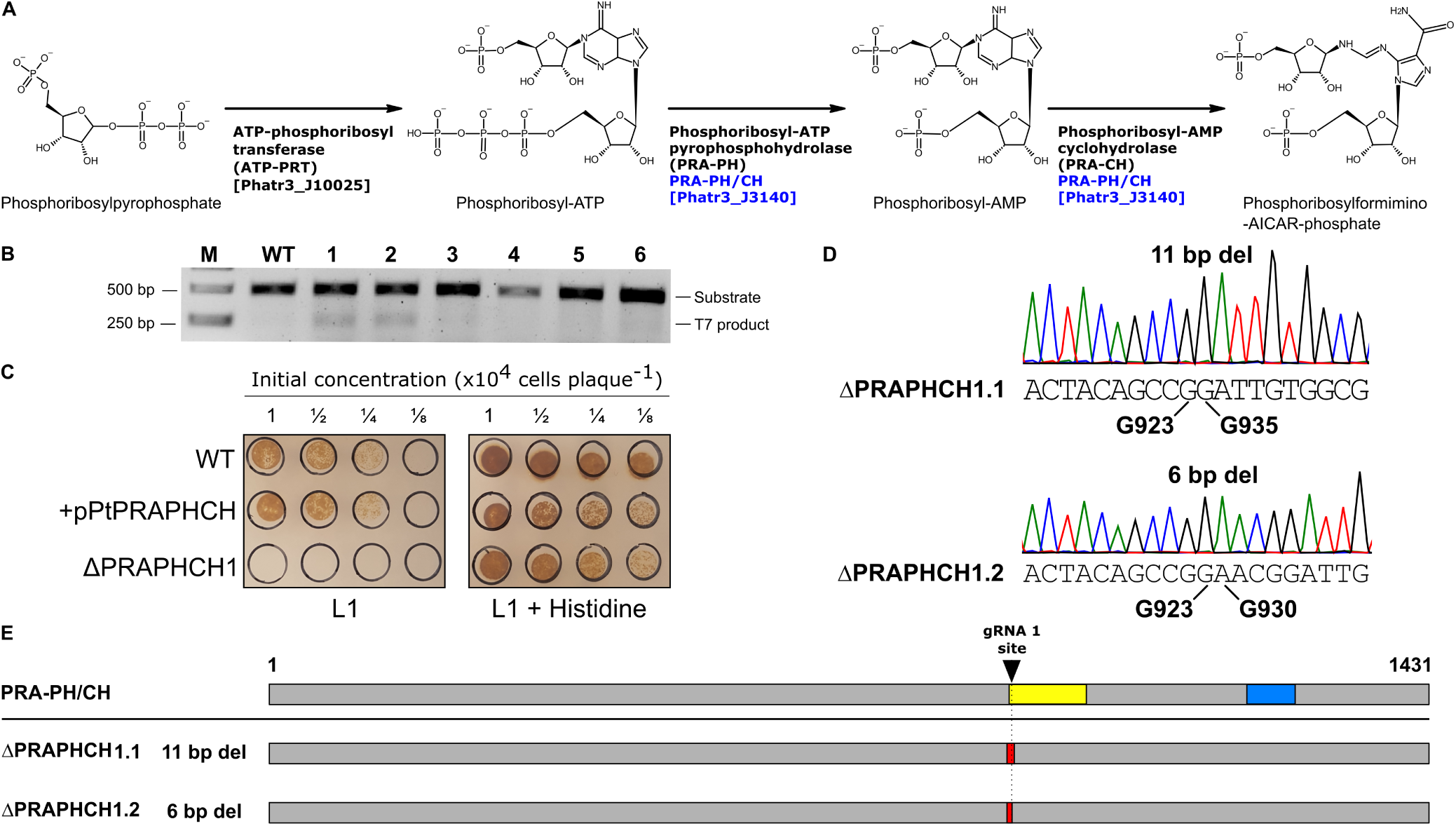
CRISPR-generated knockouts in the predicted *P. tricornutum* histidine biosynthesis pathway. (A) Predicted biosynthesis pathway for conversion of ribose-5-phosphate to L-histidine, with the bi-functional PtPRA-PH/CH enzyme highlighted in blue. Abbreviated names for each enzyme are indicated in parentheses, and the predicted corresponding *P. tricornutum* gene names are indicated in square brackets. (B) Example image of T7EI editing assay to screen exconjugants for potential editing events in the PtPRA-PH/CH gene. Substrate indicates PtPRA-PH/CH gene fragments amplified by the PCR, while T7 product indicates exconjugants with evidence of Cas9 editing. WT, wild-type *P. tricornutum* genomic DNA used in the T7EI editing assay. (C) Example of phenotypic screening of one PtPRA-PH/CH knockout strain (ΔPtPRAPHCH1) transformed with or without the complementing PRA-PH/CH plasmid on L1 solid media alone or L1 supplemented with histidine at the indicated dilution of initial concentraion. WT, wild-type *P. tricornutum* strain. (D) Sanger sequencing traces of characterized PtPRA-PH/CH knockouts with the position (below trace) and type of insertion or deletion (above trace) indicated for each allele of the three strains. (E) Graphical map of the position and extent of indels for PtPRA-PH/CH knockout relative the wild-type PtPRA-PH/CH gene (shown at top). Red rectangles indicate nucleotide deletions, while the yellow and blue rectangles on the WT gene indicated the position of the PRA-PH and CH active sites.

We also identified the PtI3GPS-PRAI gene as a potential target as it encodes a predicted bi-functional enzyme that is a fusion of indole-3-glycerol-phosphate synthase (I3PGS) and phosphoribosylanthranilate isomerase (PRAI), and would catalyze two successive steps in the tryptophan biosynthesis pathway (Supplementary Figure 3).

To confirm the genomic target sites, we PCR-amplified and sequenced the PtUMPS and PtPRA-PH/CH genes of the *P. tricornutum* CCAP 1055/1 strain used in our laboratory. Two distinct alleles for both the PtUMPS and PtPRA-PH/CH genes were identified. Seven single-nucleotide polymorphisms (SNPs) in the PtUMPS alleles result in amino acid substitutions that differentiate the two alleles from each other and from the published *P. tricornutum* genome (Supplementary Table 4). All substitutions are located in non-conserved regions of the PtUMPS protein (Supplementary Figure 4). Similarly, an A to G mutation at base position 1205 in allele 2 of the PtPRA-PH/CH gene was identified (Supplementary Table 5). This transversion converts a highly conserved glutamate to a glycine in the catalytic site of the PRA-PH domain. The impact of these substitutions on PtUMPS and PtPRA-PH/CH function is not known.

### Cas9 and TevCas9 editing of auxotrophic genes is characterized by loss of heterozygosity

To generate knockouts in uracil and histidine biosynthetic genes, we designed and individually cloned Cas9 and TevCas9 single guide RNAs (sgRNAs) against different sites in the PtUMPS and PtPRA-PH/CH genes (Table 1). The TevCas9 nuclease is a dual nuclease that generates a 33-38 base pair deletion between the I-TevI (Tev) and Cas9 cut sites^26^. The targeting requirements for a TevCas9 nuclease are an I-TevI 5*′*-CNNNG-3*′* cleavage motif positioned ∼ 15-18 base pairs upstream of the 5*′* end of the sgRNA binding site. The Cas9 or TevCas9 editing plasmids were moved into *P. tricornutum* by bacterial conjugation and exconjugants selected on Zeocin™-containing media.

**Table 1.**
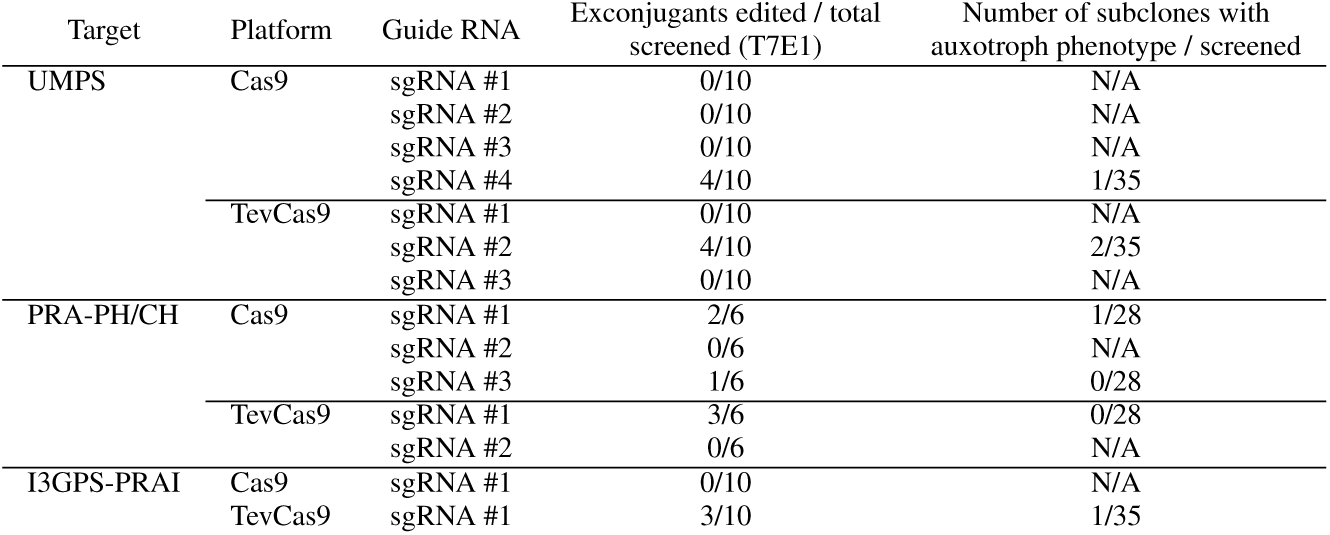
Summary of sgRNAs used for Cas9 and TevCas9 editing.

We first assessed editing by screening *P. tricornutum* exconjugants by T7 endonuclease I (T7EI) mismatch cleavage assays on PCR products amplified from each target gene ((Figure 1B and 2B, and Table 1). This assay identified 5 sgRNAs with detectable editing rates. Colonies that showed editing were diluted, plated to obtain subclones, and subsequently screened for the corresponding auxotrophic phenotype on solid media with and without auxotrophic supplement (uracil or histidine) (Figure 1C and 2C). For knockout of the PtUMPS gene, we further characterized 3 subclones with a uracil-requiring phenotype. Because the two PtUMPS alleles of *P. tricornutum* possessed SNPs relative to each other, we were able to map allele-specific editing events (Figure 1D and 1E). Two of the strains, ΔUMPS1 and ΔUMPS2, exhibited loss of heterozygosity with one allele possessing a small deletion (< 20 bps) and the other allele possessing a large deletion (> 610 bp). The third characterized subclone, ΔUMPS3, possessed a homozygous 1-bp insertion. For the PtPRA-PH/CH knockouts that generated a histidine-requiring phenotype (Figure 2B and 2C), targeted sequencing of one subclone revealed an 11-bp deletion in one allele and a 6-bp deletion in the second allele (Figure 2D and 2E).

The types of deletions observed in the uracil- and histidine-auxotrophs are consistent with heterogeneous editing events resulting in loss of heterozygosity^27–29^. To extend these observations, we used Nanopore sequencing to better assess the spectrum of large deletions that are often overlooked in Cas9-editing studies. In addition to the two sgRNAs that showed robust editing on the PtUMPS gene, we examined deletion events in exconjugants with sgRNAs targeted to the PtUREASE gene^6^ and the PtI3GPS-PRAI gene. The PtI3GPS-PRAI gene encodes a predicted bi-functional enzyme that is a fusion of indole-3-glycerol-phosphate synthase (I3PGS) and phosphoribosylanthranilate isomerase (PRAI) that is predicted to catalyze two successive steps in the tryptophan biosynthesis pathway (Supplementary Figure 3). For each experiment, ∼1000 exconjugants were pooled and a ∼6-kb PCR product generated for each of the target genes with the predicted Cas9 or TevCas9 target sites in the middle of the amplicon. We focused our attention on deletions > 50 bp as these deletions are typically under-reported in targeted amplicon sequencing (Figure 3). We noted a drop in Nanopore read coverage centered around the predicted sgRNA target sites for products amplified from Cas9 and TevCas9 editing experiments (black dots) as compared to read coverage for control experiments (orange dots), consistent with editing at those sites (Figure 3, *left* panels). Mapping the deletion start and end points revealed that most deletions were centered on the Cas9 or TevCas9 target site (Figure 3, *right* panels), with deletions extending up to 2700-bp (Figure 3, *centre* panel). The mean deletion length for all Cas9 editing events was 1735±719 bp and 2006±633 bp for TevCas9 events.

**Figure 3.**
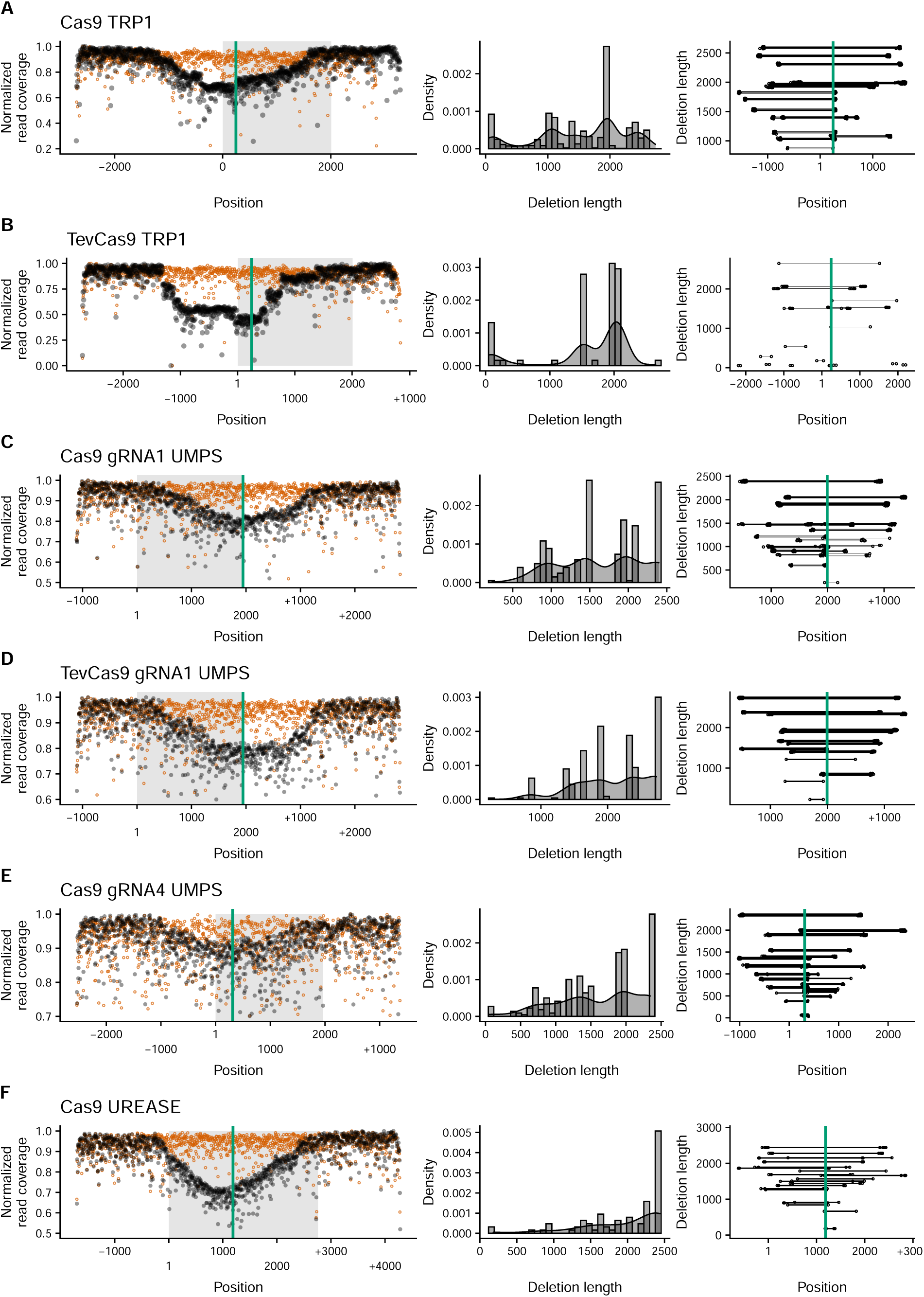
Large deletions in edited *P. tricornutum* auxotrophic genes captured by Nanopore amplicon sequencing. For each panel A-E, the name of the target gene as well as the editing enzyme are indicated. The *leftmost* plot shows normalized read coverage averaged over a 5-bp window for the edited sample (black dots) and the wild-type sample (orange dots) relative to the position in PCR amplicon. Numbering on the x-axis is relative to the ATG start codon for each gene, with sequence upstream indicated by a minus (-) symbol and sequence downstream indicated by a plus (+) symbol. The green vertical line indicates the Cas9 or TevCas9 cleavage site, while the shaded rectangle indicates the ORF. The *middle* plot is a density plot of deletions > 50-bp. The *rightmost* plot shows the length and position of deletions > 50-bp relative to their position in the PCR amplicon, with numbering of the x-axis as in the *leftmost* panel. Each horizontal line indicates a mapped deletion event. Deletions events are ordered from longest to smallest. The green line indicates the Cas9 or TevCas9 cleavage site.

Collectively, this data shows that Cas9 or TevCas9 editing of biosynthetic genes can readily generate *P. tricornutum* auxotrophs that can be identified by phenotypic or genetic screens. Moreover, our data agree with a growing body of evidence revealing that Cas9 editing (and TevCas9 editing here) generates large deletions that would typically be missed unless screening strategies are explicitly designed to look for loss of heterozygosity.

### Phenotypic and metabolomic characterization the PtUMPS knockouts

Two uracil-requiring auxotrophs (ΔUMPS1 and ΔUMPS2) were selected for further characterization by first spot plating onto L1 media with and without uracil and 5-FOA. The PtUMPS knockout strains were only able to survive in the presence of uracil supplementation. Additionally, the knockouts survived on 5-FOA concentrations that fully inhibited the growth of wild-type *P. tricornutum* (Figure 4A). This is consistent with phenotypes previously observed for *P. tricornutum* UMPS knockouts^7, 21^. There was a slight growth advantage of ΔUMPS1 over ΔUMPS2 on media supplemented with both 5-FOA and uracil, but not on media containing uracil alone. To compare if the observed phenotypes were consistent across solid and liquid media, we monitored the growth of these strains over 10 days in liquid media and found that the growth rates were consistent with those observed on solid media, with one notable difference (Figure 4B and Table 2). The growth advantage of ΔUMPS1 over ΔUMPS2 observed on solid media supplemented with both 5-FOA and uracil was not replicated in liquid media as the generation times for ΔUMPS1 and ΔUMPS2 were very similar (∼24 and ∼22 hours, respectively).

**Figure 4.**
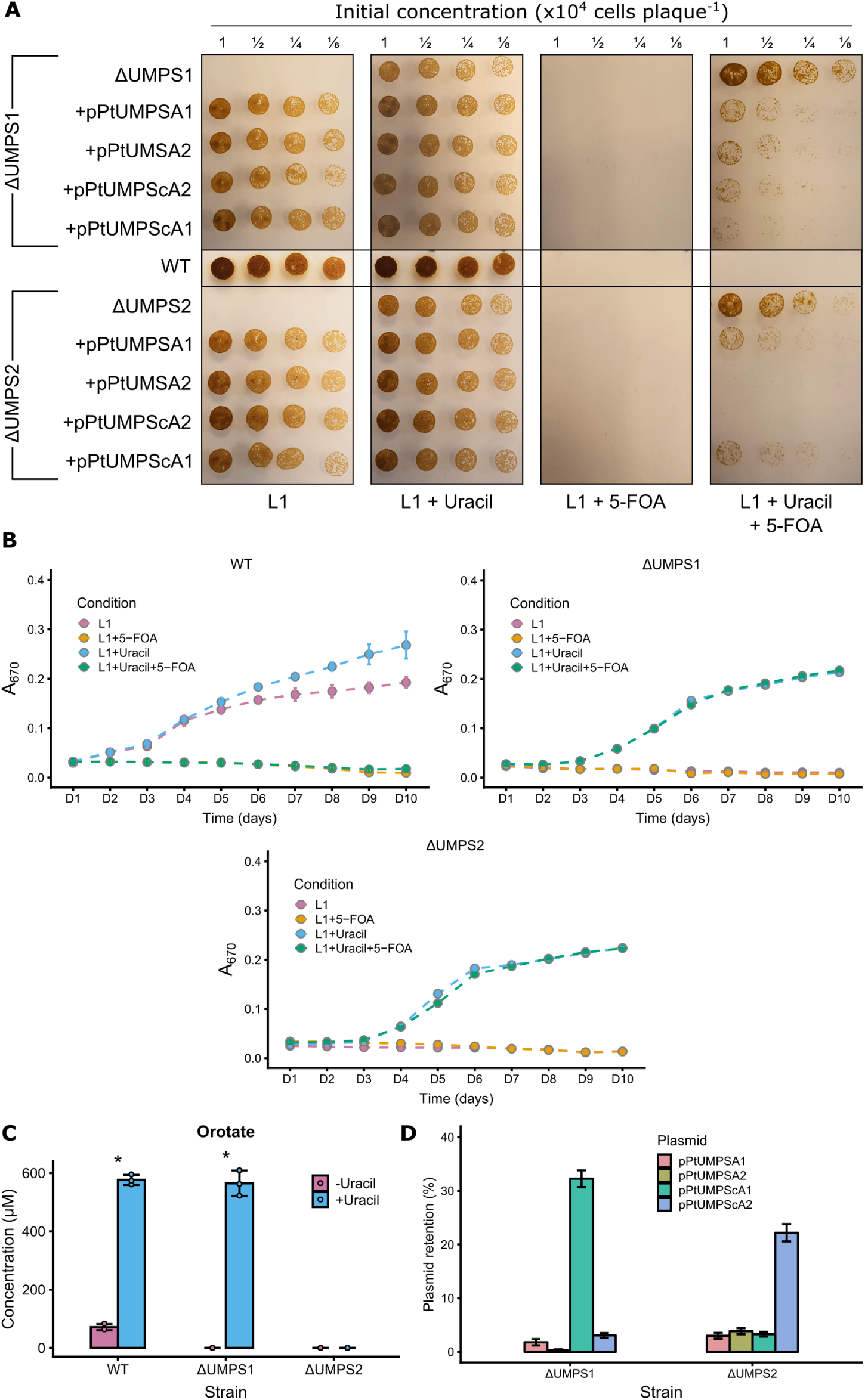
Phenotypic and metabolomic characterization of PtUMPS knockouts. (A) Spot plating assays of wild type (WT), ΔUMPS1 over ΔUMPS2 strains on L1 solid media alone, or L1 supplemented with uracil or 5-FOA, or both. Indicated dilutions are relative to the initial concentration. (B) Liquid growth curves of wild type (WT), ΔUMPS1 over ΔUMPS2 strains in L1 liquid media alone, or supplemented with uracil or 5-FOA or both. Data points are the mean of three independent replicates, with error bars representing the standard error of the mean. (C) Orotate concentrations were measured by LC-MS for cultures grown with and without uracil supplementation. Bars represent mean values and error bars represent standard deviation for three biological replicates. Individual data points are represented as colored dots. Statistical confidence level was calculated by one-sided t test. p < 0.001 is indicated by an asterisk. (D) Bar graph showing percent plasmid retention in the ΔUMPS1 and ΔUMPS2 strains harbouring various PtUMPS constructs after 14 days of outgrowth. Bars represent the mean ratio of colonies on selective L1 + nourseothricin versus non-selective L1 plates from three independent replicates, with error bars representing the standard error of the mean.

To investigate the impact of PtUMPS knockouts on uracil metabolism, we performed targeted metabolomics on the UMPS substrate orotate using LC-MS in wild-type and knockouts strains (Figure 4C). We focused characterizing the orotate intermediate in the uracil pathway (Figure 2A) predicting that there should be an increase of orotate in knockout strains relative to wild type. We were unable to detect orotate in the ΔUMPS1 strain in the absence of uracil supplementation (-uracil), or in the ΔUMPS2 strain in either the -uracil or +uracil condition. A ∼6-fold increase of cellular orotate levels was observed in the wild-type strain when L1 media was supplemented with uracil (+uracil) as compared to minimal L1 media (-uracil) (Figure 4C). Interestingly, when the ΔUMPS1 strain was grown with uracil supplementation we detected orotate at levels similar to those observed in the wild-type strain grown with uracil. This result suggests that allele 1 in the ΔUMPS1 knockout strain (with an 18-bp in-frame deletion) retains UMPS activity that behaves similar to the wild-type strain. In contrast, the ΔUMPS2 strain has two out-of-frame deletions that likely abolish UMPS and OPRT activity. We speculate that undetectable levels of orotate may be because it is diverted to another biosynthetic pathway in the ΔUMPS2 strain.

### Plasmid complementation of the uracil and histidine auxotrophs

Plasmid-based complementation of *P. tricornutum* auxotrophs would validate that the Cas9-editing event was the cause of the auxotrophic phenotype, as well as providing alternatives to antibiotic-based selection methods to maintain episomal vectors. We first examined complementation of the uracil-requiring phenotype by cloning both gDNA and cDNA versions of each PtUMPS allele with the native promoter and terminator into the nourseothricin-resistant pPtGE31 expression plasmid^6^ (Supplementary Dataset 1). These plasmids were designated pPtUMPSA1, pPtUMPSA2, pPtUMPScA1, and pPtUMPScA2 (Supplementary Table 2) and moved into the ΔUMPS1 and ΔUMPS2 strains via conjugation. Exconjugants were spot-plated onto solid L1 media with and without uracil and 5-FOA supplementation (Figure 4A). All complemented strains grew on minimal L1 media, while the uncomplemented knockouts did not, confirming expression of the UMPS gene from the pPtGE31 plasmid. No strain grew on 5-FOA alone. Unexpectedly, some of the complemented strains survived on plates supplemented with both 5-FOA and uracil. For example, when ΔUMPS2 was transformed with either of the allele 1 complementation plasmids (pPtUMPSA1 and pPtUMPScA1), clear resistance to 5-FOA in the presence of uracil was observed. The phenotypes observed on solid media were consistent with those observed when the strains were grown in liquid media with similar media supplementation.

The growth phenotype of the ΔUMPS1 and ΔUMPS2 strains in media supplemented with uracil and 5-FOA could be explained by counter-selection against the plasmid carrying an intact PtUMPS gene that would metabolize 5-FOA to a toxic intermediate. We thus tested for plasmid loss in the complemented strains by plating the ΔUMPS1 and ΔUMPS2 strains carrying different expression plasmids on solid L1 + nourseothricin after 14 days of growth. As shown in Figure 4D, plasmid retention in all strains was severely reduced, ranging from ∼1% to ∼33%, explaining why colonies readily appeared on L1 media supplemented with 5-FOA and uracil. These observations suggest that curing of plasmids carrying the PtUMPS gene from PtUMPS knockout strains is a simple matter of growth on the appropriate media.

Similarly, we were able to complement the histidine-requiring phenotype by cloning a wild-type copy of the PtPRA-PH/CH gene into an expression vector, and transforming the plasmid into the ΔPRAPHCH1 strain by conjugation. The ΔPRAPHCH1 strain with the complementing plasmid grew on both L1 solid with and without histidine supplementation, whereas the ΔPRAPHCH1 strain without the complementing plasmid only grew on L1 media with histidine supplementation (Figure 2C).

### Implications for expanding the *P. tricornutum* genetic toolbox

The available tools for genetic manipulation of *P. tricornutum* have grown substantially in recent years, including the adaptation of TALEN and Cas9 genome-editing nucleases for targeted knockouts^6–9, 30^. Examination of edited sites in *P. tricornutum* and other organisms has revealed that many Cas9 events are not homogeneous. Indeed, Nanopore long-read sequencing of PCR amplicons indicates that many Cas9 and TevCas9 editing events result in deletions up to ∼2.7-kb in length. Cas9 editing events that create large deletions could be a relatively straightforward approach to delete entire genes, regulatory regions, or functional domains in *P. tricornutum*.

The creation of auxotrophic strains of *P. tricornutum* with plasmid based rather than chromosomally integrated complementation markers is critical for a number of reasons. Auxotrophic strains expand the available selection schemes beyond traditional antibiotic markers and provide a facile method for strain cataloging and validation. Antibiotic-free selection is also an advantage when *P. tricornutum* is used for production of human therapeutics. In the case of uracil auxotrophs, complementing plasmids can be cured (or counter selected) by simple inclusion of 5-FOA and uracil in the growth media. We have previously shown that plasmids are lost from *P. tricornutum* by passaging cultures over multiple days in the absence of antibiotic selection required for maintenance of the plasmid^31^. However, the counter selection method by 5-FOA and uracil supplementation is more rapid and requires screening significantly fewer colonies to confirm plasmid loss. The ability to rapidly cure plasmids will be of tremendous value to prevent prolonged expression of Cas9 and possible toxicity issues during strain engineering, to cure incompatible plasmids, or to cure reporter or expression plasmids under distinct growth conditions. We also envision that rapid curing of plasmids would allow recycling of a limited number of selection markers for serial transformations needed for strain construction or genomic engineering.

## Methods

### Microbial strains and growth conditions

*Saccharomyces cerevisiae* VL6-48 (ATCC MYA-3666: *MATα his3-δ 200 trp1-δ 1 ura3-52 lys2 ade2-1 met14* °) was grown in rich medium (YPD) or complete minimal medium lacking histidine (Teknova) supplemented with 60 mg L^-1^ adenine sulfate. Complete minimal media used for spheroplast transformation contained 1 M sorbitol. *Escherichia coli* (Epi300, Epicenter) was grown in Luria Broth (LB) supplemented with appropriate antibiotics (chloramphenicol 25 mg L^-1^ or kanamycin 50 mg L^-1^ or ampicillin 50 mg L^-1^ or gentamicin 20 mg L^-1^). *Phaeodactylum tricornutum* (Culture Collection of Algae and Protozoa CCAP 1055/1) was grown in L1 medium without silica, with or without uracil (50 mg L^-1^) or histidine (200 mg L^-1^) or 5-FOA (100 mg L^-1^), supplemented with appropriate antibiotics Zeocin™ (50 mg L^-1^) or nourseothricin (100 mg L^-1^), at 18 °C under cool white fluorescent lights (75 *µ*E m^-2^ s^-1^) and a photoperiod of 16 h light:8 h dark. L1 media supplemented with nourseothricin contained half the normal amount of aquil salts. *P. tricornutum* auxotroph genotypes are as follows. Mutations in PtUMPS are described in reference to the chromosome 6 sequence (GenBank: CM000609.1), and mutations for PtPRA-PH/CH are in reference to the chromosome 3 sequence (GenBank: CP001142.1). Mutations described for each gene are listed for allele 1 followed by allele 2, and numbered beginning from the first nucleotide of the start codon for simplicity. Genotypes of auxotroph strains generated in this study are listed in Supplementary Table 1.

### Transfer of DNA to P. tricornutum via conjugation from E. coli

Conjugations were performed as previously described^5, 6^. Briefly, liquid cultures (250 *µ*l) of *P. tricornutum* were adjusted to a density of 1.0 × 10^8^ cells mL^-1^ using counts from a hemocytometer, plated on 1/2 x L1 1% agar plates and grown for four days. L1 media (1.5 mL) was added to the plate and cells were scraped and the concentration was adjusted to 5.0 × 10^8^ cells mL^-1^. *E. coli* cultures (50 mL) were grown at 37°C to A600 of 0.8–1.0, centrifuged for 10 min at 3,000 x g and resuspended in 500 *µ*l of SOC media. Conjugation was initiated by mixing 200 *µ*l of *P. tricornutum* and 200 *µ*l of *E. coli* cells. The cell mixture was plated on 1/2 x L1 5% LB 1% agar plates, incubated for 90 minutes at 30°C in the dark, and then moved to 18°C in the light and grown for 2 days. After 2 days, L1 media (1.5 mL) was added to the plates, the cells scraped, and 300 *µ*l (20%) plated on 1/2 X L1 1% agar plates supplemented with Zeocin™ 50 mg L^-1^ or nourseothricin 200 mg L^-1^. Colonies appeared after 7–14 days incubation at 18°C with light.

### Plasmid design and construction

All plasmids were constructed using a modified yeast assembly protocol^32, 33^. Plasmids pPtUMPSA1 and pPtUMPSA2 were made from pPtGE31^6^ by replacing the URA3 element with a PCR fragment consisting of PtUMPS allele 1 or 2 with ∼1 kb up- and downstream of the PtUMPS ORF, amplified from *P. tricornutum* genomic DNA. Plasmids pPtUMPScA1 and pPtUMPScA2 were made from pPtUMPSA1 and pPtUMPSA2 by replacing the PtUMPS ORF with a PCR fragment consisting of PtUMPS allele 1 or 2 amplified from *P. tricornutum* cDNA. Plasmid pPtUMPS40S was made from pPtGE31 by replacing the URA3 element with a cassette consisting of PCR fragments of the 40SRPS8 promoter and terminator^6^ flanking a PCR fragment of the PtUMPS allele 1 ORF amplified from *P. tricornutum* genomic DNA. Plasmid pPtPRAPHCH was made from pPtGE31^6^ by replacing the URA3 element with a PCR fragment consisting of PtPRA-PH/CH with ∼1 kb up- and downstream of the PtPRA-PH/CH ORF, amplified from *P. tricornutum* genomic DNA. Using golden gate assembly, sgRNAs targeting different regions of the PtUMPS and PtPRA-PH/CH genes were cloned into the BsaI sites positioned between the *P. tricornutum* U6 promoter and terminator in pPtGE34 and pPtGE35. Plasmid constructs were confirmed by Sanger sequencing at the London Regional Genomics Facility.

### Generation of PtUMPS and PtPRA-PH/CH knockouts using Cas9 and TevCas9

Plasmids pPtGE34 or pPtGE35, containing no guide RNA or sgRNA#1, sgRNA#2, sgRNA#3 or sgRNA#4 for the PtUMPS gene, or sgRNA#1 or sgRNA#2 for the PtPRA-PH/CH gene, were conjugated from *E. coli* to *P. tricornutum* and exconjugants were selected on Zeocin™-containing media, supplemented with uracil or histidine as appropriate^34^. Ten colonies from each conjugation were resuspended in TE buffer and flash frozen at −80°C followed by heating at 95°C to lyse cells and extract genomic DNA. The genomic target site of each sgRNA in *P. tricornutum* was amplified by PCR and the products were analyzed by T7EI assay as follows; PCR products were denatured at 95°C for 5 minutes, slowly cooled to 50°C, and flash frozen at −20°C for 2 minutes. PCR products (250 ng) were incubated with 2U of T7EI (NEB) in 1x NEBuffer 2 for 15 min at 37°C and analyzed by agarose gel electrophoresis. Colonies that showed editing by T7EI assay were grown in liquid culture supplemented with Zeocin™ and uracil or histidine as appropriate for 2 weeks and serial dilutions were plated on selective media with uracil or histidine to isolate sub-clones. Sub-clones were then screened for homozygous PtUMPS or PtPRA-PH/CH knockout phenotypes by replica streaking on minimal L1 media and L1 media supplemented with uracil or histidine as appropriate. Streaks were grown for 5 days before visual identification of phenotypes. Sub-clones that were identified as phenotypic knockouts were resuspended in TE buffer and flash frozen at −80°C followed by heating at 9°C to lyse cells and extract genomic DNA, then sgRNA target sites were PCR amplified. Sanger sequencing of PCR products was performed at the London Regional Genomics Facility to identify the type and length of indels generated. Stable bi-allelic PtUMPS or PtPRA-PH/CH knockout mutant lines were then grown in nonselective L1 media supplemented with uracil or histidine for 1 week to cure them of plasmids before plating to obtain single colonies. Resulting colonies were replica streaked onto nonselective and Zeocin™-containing media supplemented with uracil or histidine to identify colonies which had successfully been cured of the plasmid.

### Spot plating P. tricornutum

Cultures of *P. tricornutum* were adjusted to 1 × 10^6^ cells mL^-1^ and serially diluted 2 X three times. For uracil auxotrophs, 10 *µ*L of each adjusted culture and dilutions were spot plated onto minimal L1 media and L1 media supplemented with uracil (50 mg L^-1^), 5-FOA (100 mg L^-1^), or both. For histidine auxotrophs, 10 *µ*L of each adjusted culture and dilutions were spot plated onto minimal L1 media and L1 media supplemented with histidine (200 mg L^-1^). Plates were incubated at 18°C under cool white fluorescent lights (75 *µ*E m^-2^ s^-1^) and a photoperiod of 16 h light:8 h dark for 7-10 days.

### Measuring P. tricornutum growth rates

Growth was measured in a Multiskan Go microplate spectrophotometer. Cultures of each strain (WT, ΔUMPS1, ΔUMPS1 + pPtUMPS40S, ΔUMPS1 + pPtUMPSA1, ΔUMPS1 + pPtUMPSA2, ΔUMPS1 + pPtUMPScA1, ΔUMPS1 + pPtUMPScA2, ΔUMPS2, ΔUMPS2 + pPtUMPSA1, ΔUMPS2 + pPtUMPSA2, ΔUMPS2 + pPtUMPScA1, ΔUMPS2 + pPtUMPScA2) were adjusted to 5 × 10^5^ cells mL^-1^ in L1 media with and without supplemented uracil (50 mg L^-1^), 5-FOA (100 mg L^-1^), or both. Two hundred microliters of each adjusted culture was added to three wells (technical replicates) of a 96-well microplate. The 96-well microplates were incubated at 18°C under cool white fluorescent lights (75 *µ*E m^-2^ s^-1^) and a photoperiod of 16 h light:8 h dark for 10 days, and absorbance at 670 nm (A_670_) was measured every 24 hours. The 96-well microplates were shaken briefly to resuspend any settled cells prior to absorbance measurements. Note that the A_670_ was not adjusted for path length and light scattering from the microplate lid and is therefore not directly comparable to optical density readings measured in a standard cuvette.

### P. tricornutum metabolite extraction

Cultures of *P. tricornutum* (Wild-type, ΔUMPS1, and ΔUMPS2) were grown with and without uracil supplementation and harvested during exponential phase as follows (Note: The ΔUMPS1 and ΔUMPS2 cultures were first grown with uracil supplementation, then switched to minimal L1 media for 1 week prior to harvesting). Cultures (∼1×10^9^ cells) were pelleted by centrifugation at 4000 × g for 10 minutes and washed by resuspending in fresh L1 media. Cells were pelleted again, resuspended in a small volume (∼5 mL) of L1 media, and transferred to a clean 10 mL syringe (without needle) with the exit plugged by parafilm. The syringe was placed, tip-down, into a clean 50 mL falcon tube and the cells were pelleted as above. The supernatant was removed and the pellet was slowly ejected from the syringe into a pre-chilled mortar containing liquid nitrogen. The frozen cells were ground to a fine powder and then transferred to a clean pre-weighed 1.5 mL eppendorf tube, suspended half way in liquid nitrogen. Being careful to keep samples frozen, 50 mg of frozen ground powder was weighed out into a new clean 1.5 mL eppendorf tube, pre-cooled in liquid nitrogen, and 250 *µ*L of cold extraction buffer with internal standard (IS) (80% methanol in MilliQ water, 125 *µ*M ^15^N_2_-uracil) was added. Samples were then homogenized by vigorous vortexing for ∼30 seconds in 10 second intervals, between which samples are kept on ice for ~30 seconds. Homogenized samples were then spun down at 4°C for 10 minutes at 20,000 x g. The supernatant was transferred to a new clean 1.5 mL eppendorf tube and spun down at 20,000 x g for 5 minutes at 4°C. The supernatant was again transferred to a new clean 1.5 mL eppendorf tube and kept at 4°C overnight until LC-MS analysis. The IS was added to the samples to compensate for losses that might occur during preparation of the samples and loss of sensitivity attributable to quenching of the signal by coeluting compounds.

### Chromatographic separation and mass spectrometry

Metabolites were separated at 45°C on a Waters Acquity HSS T3 column [2.1 × 100 mm, 1.8 *µ*m particle] in a Waters ACQUITY UPLC I-Class system (Waters, Milford, MA). Solvent A consisted of water and solvent B consisted of methanol, both containing 0.1% formic acid. Elution was performed by use of a linear gradient, at a flow rate of 0.3 mL/min, as follows: 0–2 min, 100% solvent A to 90% solvent B; 2.01 minutes, 100% solvent A to recondition the column. A Waters Xevo G2-S quadrupole time of flight mass spectrometer was operated in negative electrospray ionization (ESI) in resolution mode. The capillary voltage was set to 1.0 kV, the source temperature was 150°C, desolvation temperature was 600°C, the cone gas was 50 L/hr and the desolvation gas was 1000 L/hr. Leucine enkephalin was infused as the lock mass with a scan time of 0.3 seconds every 10 seconds and three scans were averaged. Linearity and detection limits for each compound were established by injection of calibration mixtures with different concentrations (0, 1, 2, 4, 8, 16, 31.25, 62.5, 125, 250, and 500 *µ*mol/L). Stable-isotope-labeled uracil (^15^N_2_-uracil) was used as the IS. The concentration of each analyte was determined by use of the slope and intercept of the calibration curve that was obtained from a least-squares regression for the analyte/IS peak-area ratio vs the concentration of the analyte in the calibration mixture.

### P. tricornutum DNA extraction and targeted long-read sequencing

Plasmids pPtGE34 or pPtGE35, containing sgRNAs targeting the PtUMPS, PtUrease, or PtI3GPS-PRAI gene were conjugated from *E. coli* to *P. tricornutum* and exconjugants were selected on Zeocin™-containing media, supplemented with uracil or tryptophan (100 mg L^-1^) as appropriate. For each transformation, colonies (∼1000) were scraped and pooled in liquid L1 media and genomic DNA was extracted using a modified akaline lysis protocol as follows: Cells were pelleted at 4000g for 5 min, and resuspended in 250 *µ*L resuspension buffer consisting of 235 *µ*L P1 (Qiagen), 5 *µ*L hemicellulose 100 mg mL^-1^, 5 *µ*L of lysozyme 25 mg mL^-1^, and 5 *µ*L zymolyase solution (200 mg zymolyase 20 T (USB), 9 mL H2O, 1 mL 1 M Tris pH 7.5, 10 mL 50% glycerol) and incubated at 37 °C for 30 min. Next, 250 *µ*L of lysis buffer P2 (Qiagen) was added, followed by 250 *µ*L of neutralization buffer P3 (Qiagen) and centrifugation at 16000g for 10 min. The supernatant was transferred to a clean tube, 750 *µ*L isopropanol was added, and the samples centrifuged at 16000g for 10 min. A 70% EtOH wash was performed, centrifuged at 16000g for 5 min, and pellets briefly dried, resuspended in 50–100 *µ*L of TE buffer, and incubated at 37°C for 30–60 min to dissolve.

The sgRNA target site regions were PCR amplified from sgRNA transformant genomic DNA samples, as well as a wild-type sample, with PrimeStar GXL polymerase (Takara) using primers positioned ∼3 kb up- and downstream of the target site (Table 3). PCR products were purified and DNA libraries were prepared, barcoded, and pooled using an Oxford Nanopore Ligation Sequencing Kit (SQK LSK109) and Native Barcoding Expansion 1-12 (EXP-NBD104) kit according to manufacturers protocol with the follow modification. The end prep incubation time was extended to 15 minutes at 20°C. The pooled library was then loaded on to a MinION R9.4.1 flowcell and sequenced.

### Targeted long-read sequencing analysis

After sequencing on an R9.4.1 flowcell, base calling was performed using GPU Guppy with the high accuracy configuration file version 3.4.4 (https://community.nanoporetech.com). Reads in each barcode were filtered using NanoFilt^35^ for a minimum average read quality score of 10 and a minimum read length of 2000, mapped using minimap2^36^ and filtered for reads that map to within 100 bases of each end of the reference sequence (the unedited 6kb PCR product sequence) to remove short fragments. The filtered reads were mapped using minimap2 (parameters: -ax map-ont) and outputted in sam format, then converted to bam, sorted, and indexed using samtools^37^. The per-base coverage depth for each barcode was calculated using Mosdepth^38^. All plots were created in R using the ggplot2 package^39^.

## Acknowledgements

This work was supported by Natural Sciences and Engineering Research Council of Canada (NSERC) Discovery Grants [RPGIN-2015-04800 to D.R.E. and RGPIN-2018-06172 to B.J.K.].

## Author contributions statement

S.S.S., D.J.G.,B.L.U., B.J.K. and D.R.E. conceived the experiments, S.S.S., H.W., C.K., D.J.G., and B.L.U. performed the experiments, S.S.S, D.J.G., B.J.K. and D.R.E. analysed the results, S.S.S and D.R.E. wrote the manuscript. All authors reviewed the manuscript.

## Additional information

The cDNA sequences of PtUMPS and PtPRA-PH/CH alleles have been deposited to GenBank under accession codes **MN242208, MN242209, MN242210**, and **MN242211**.

## Supplemental Information

**Figure 1.**
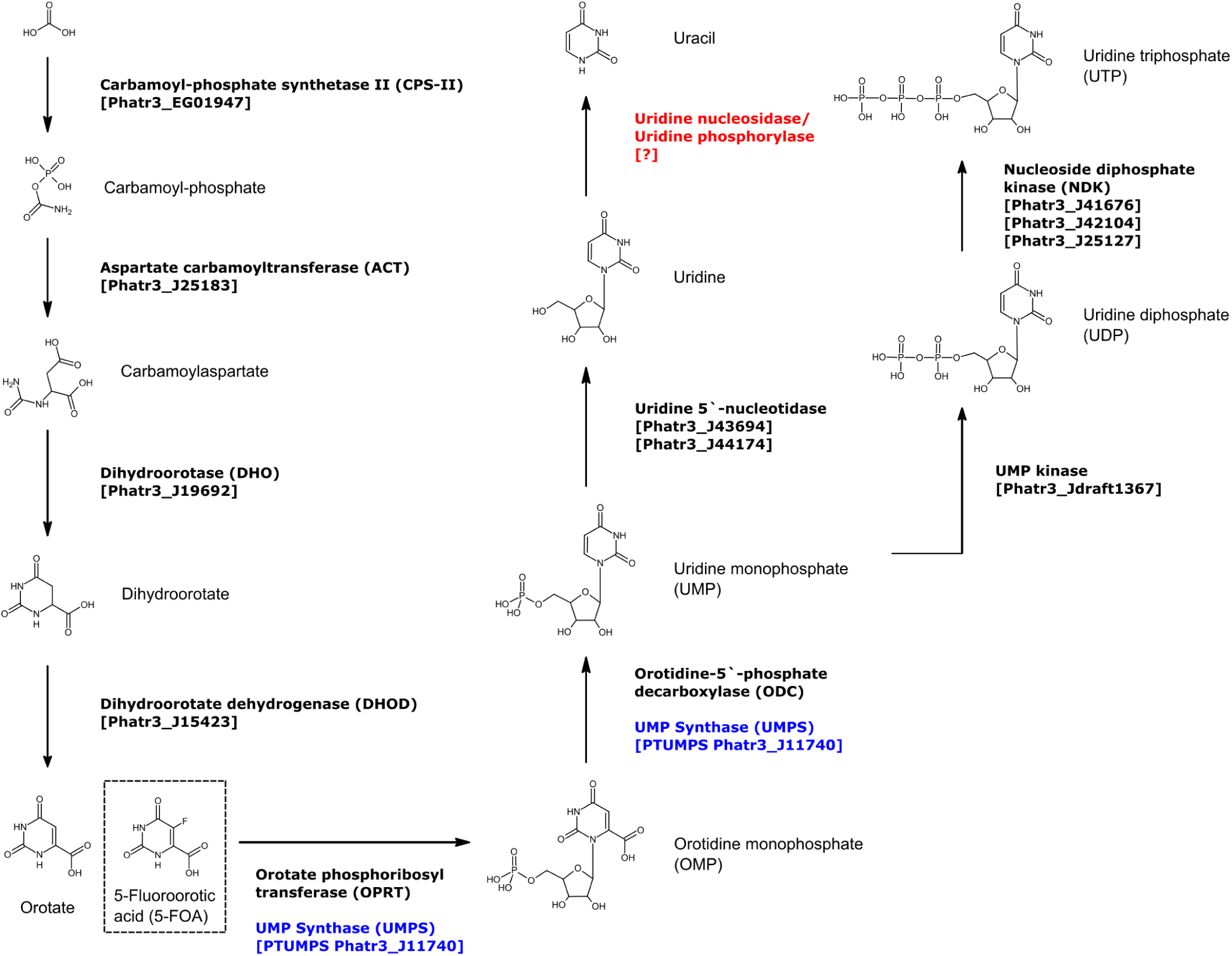
The predicted P. tricornutum uracil biosynthesis pathway. The diagram depicts the biosynthesis pathway for conversion of carbonic acid to uracil and uridine triphosphate, with the PtUMPS enzyme highlighted in blue. The negative selector, 5-Fluoroorotic acid (5-FOA), is also shown in a hashed box at the position where it enters the pathway. No enzymes sharing significant homology with uridine nucleosidase or uridine phosphorylase (highlighted in red) were identified in *P. tricornutum* by NCBI and EnsemblProtists BLAST queries. Abbreviated names for molecules and enzymes are indicated in parentheses, and the predicted corresponding *P. tricornutum* gene names are indicated in square brackets.

**Figure 2.**
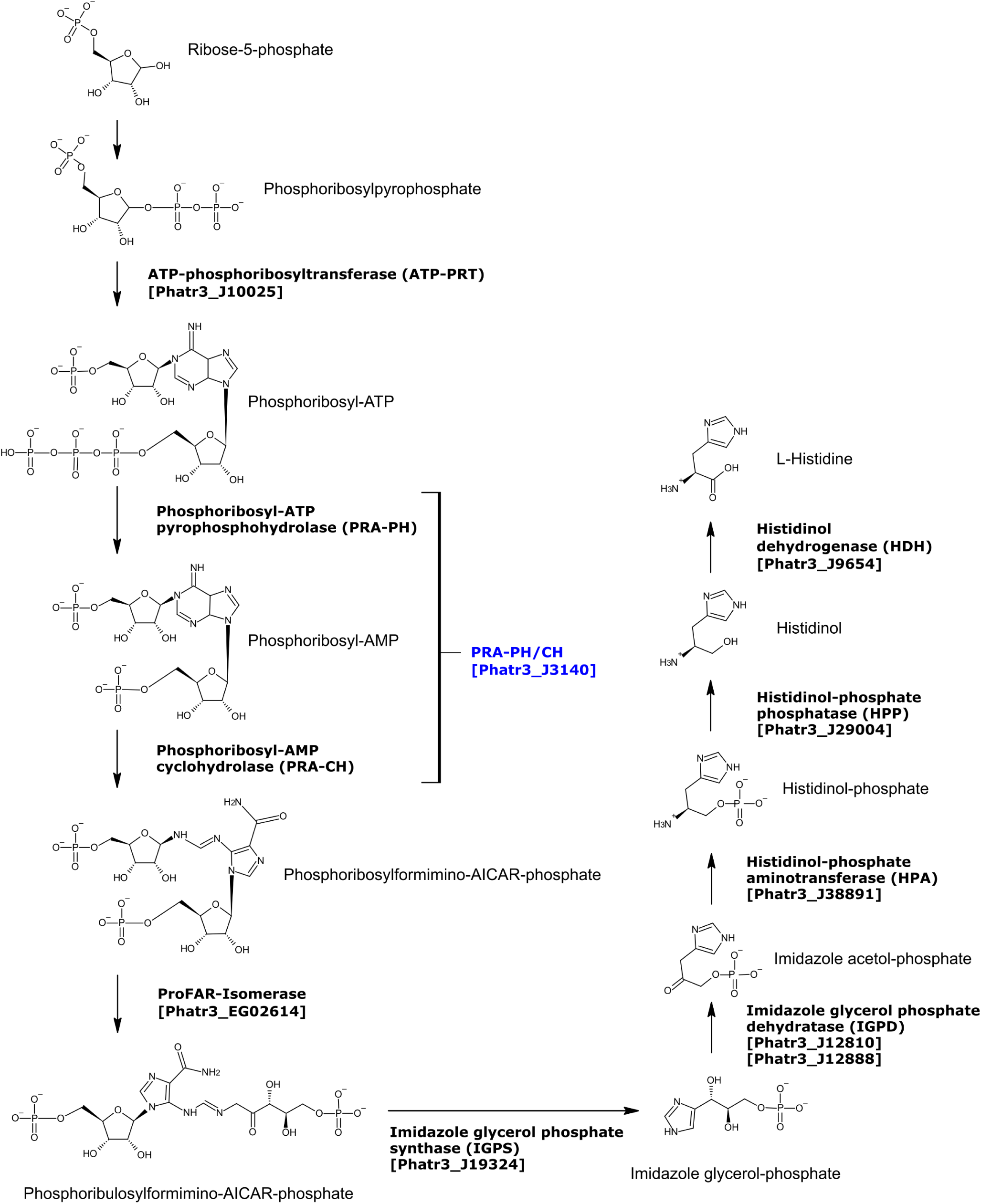
The predicted *P. tricornutum* histidine biosynthesis pathway. The diagram depicts the biosynthesis pathway for conversion of ribose-5-phosphate to L-histidine, with the PtPRA-PH/CH enzyme highlighted in blue. No enzymes sharing significant homology with HPP (highlighted in red) were identified in *P. tricornutum* by NCBI and EnsemblProtists BLAST queries. Abbreviated names for each enzyme are indicated in parentheses, and the predicted corresponding *P. tricornutum* gene names are indicated in square brackets.

**Figure 3.**
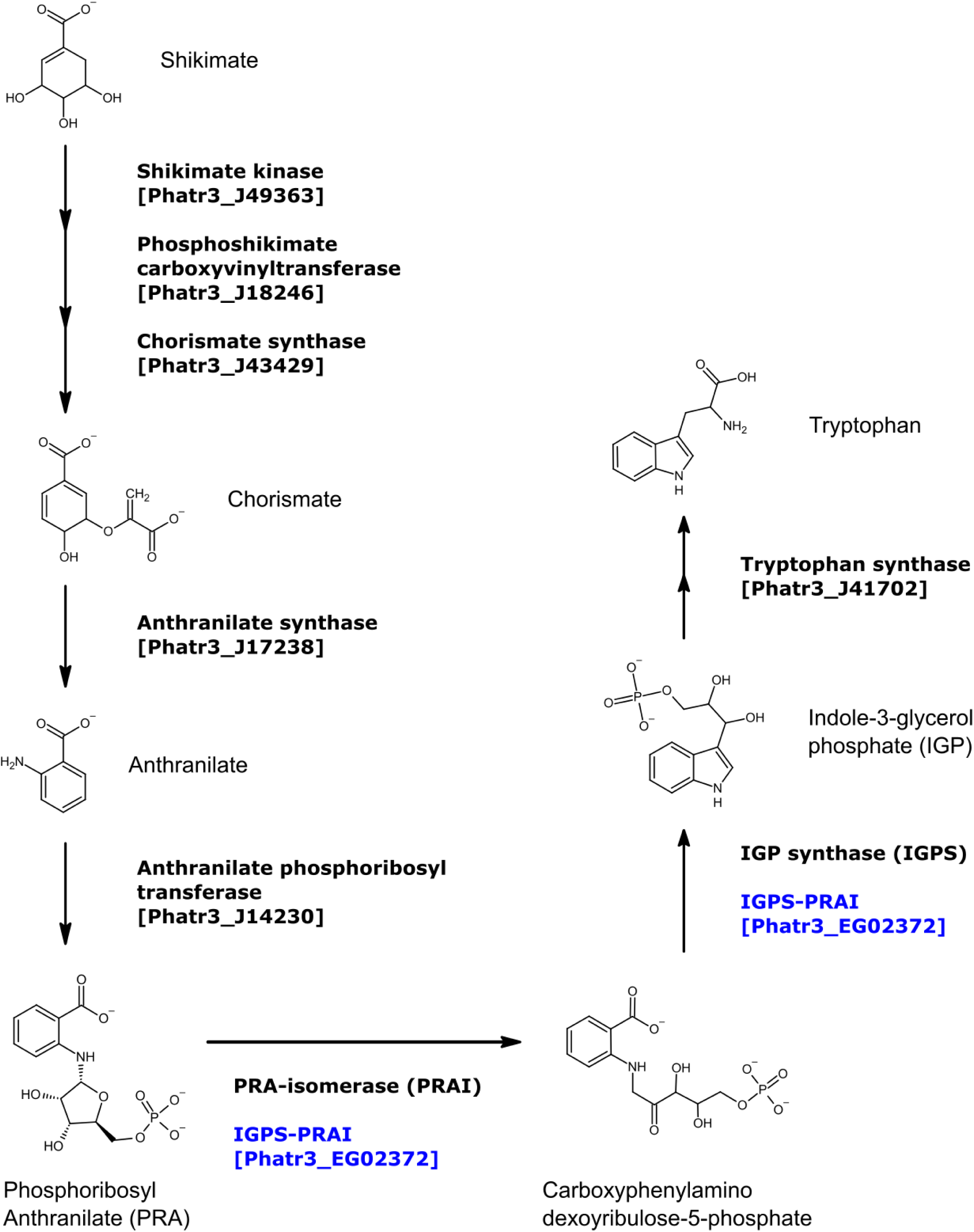
The PtI3GPS-PRAI coding sequence and gRNA target sites. The coding regions for the I3GPS (blue) and PRAI (yellow) catalytic residues are highlighted. Residues that differ between the two alleles are indicated by red boxes.

**Figure 4.**
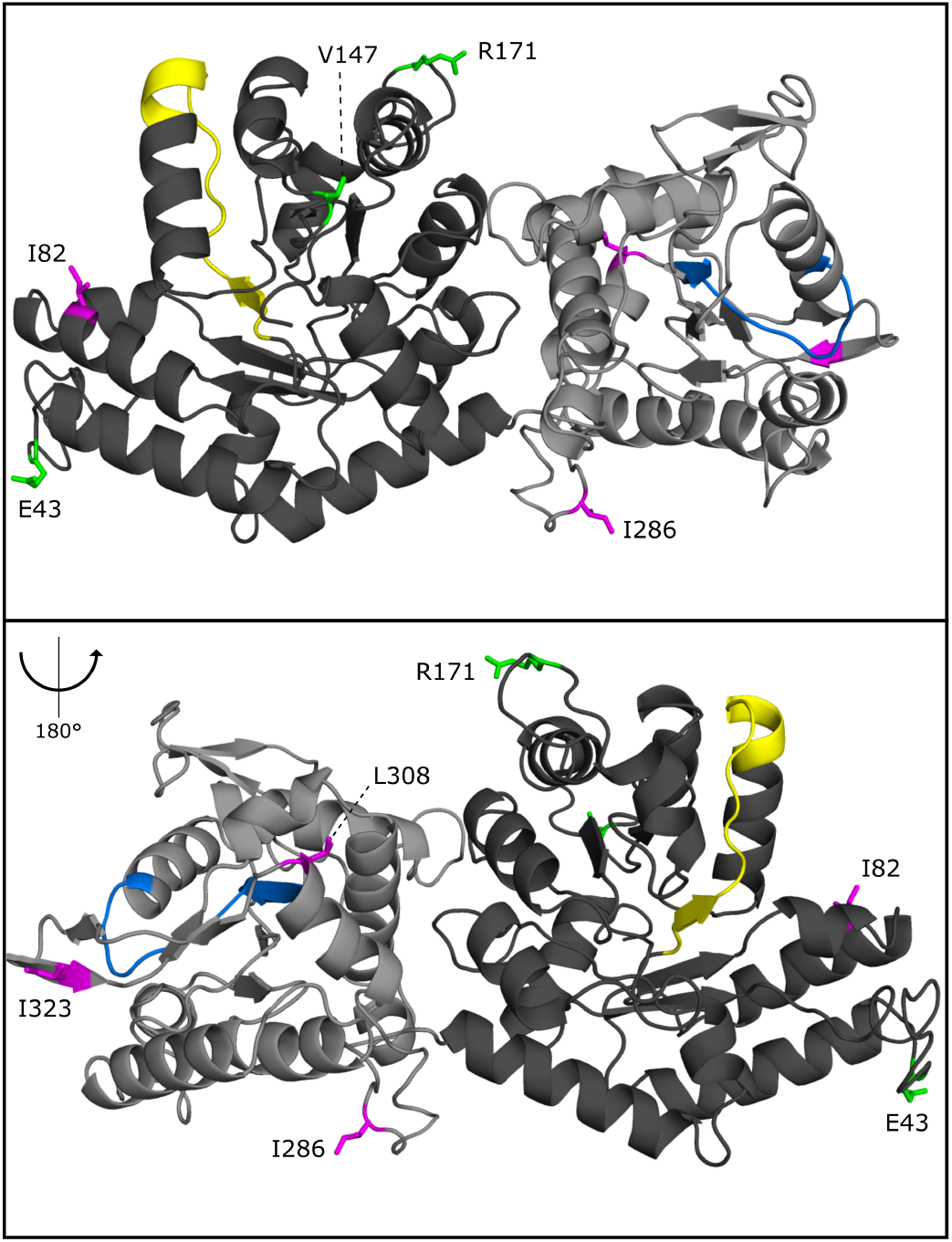
Predicted PtUMPS structure indicating the positions of the ODC (dark grey) and OPRT (light grey) domains. Regions containing conserved active site residues for the ODC and OPRT domains are indicated in yellow and blue, respectively. Residue substitutions that differentiate the two alleles are labeled and highlighted in green (allele 1) and magenta (allele 2). Folding prediction was modeled using the PHYRE2 Protein Fold Recognition Server.

**Figure 5.**
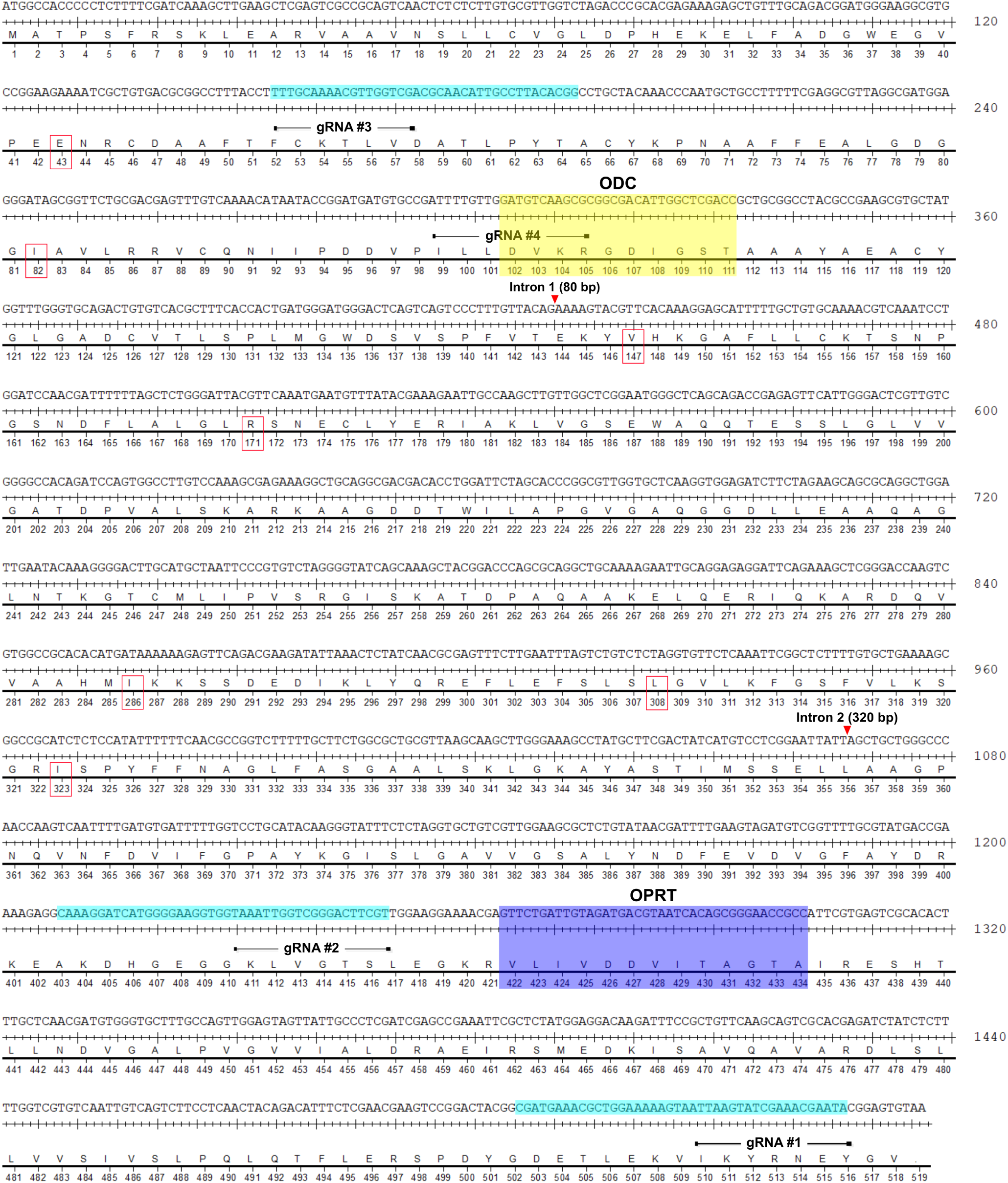
The PtUMPS coding sequence and gRNA target sites. The coding regions for the ODC (yellow) and OPRT (blue) catalytic residues, and TevCas9 target sites (light blue) are highlighted. Residues that differ between the two alleles are indicated by red boxes.

**Figure 6.**
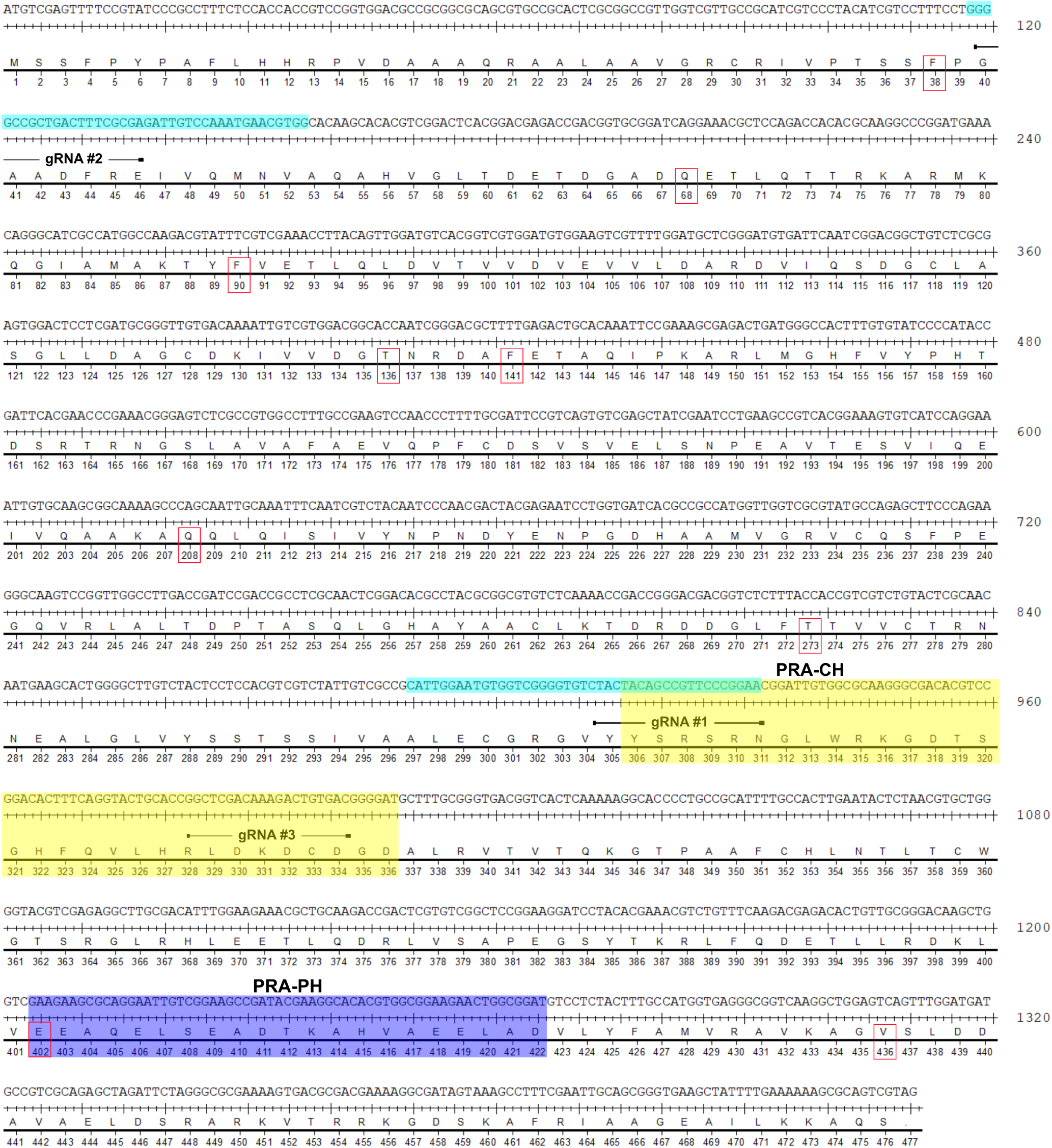
The PtPRA-PH/CH coding sequence and gRNA target sites. The coding regions for the PRA-PH (blue) and PRA-CH (yellow) catalytic residues, and TevCas9 target sites (light blue) are highlighted. Residues that differ between the two alleles are indicated by red boxes.

**Figure 7.**
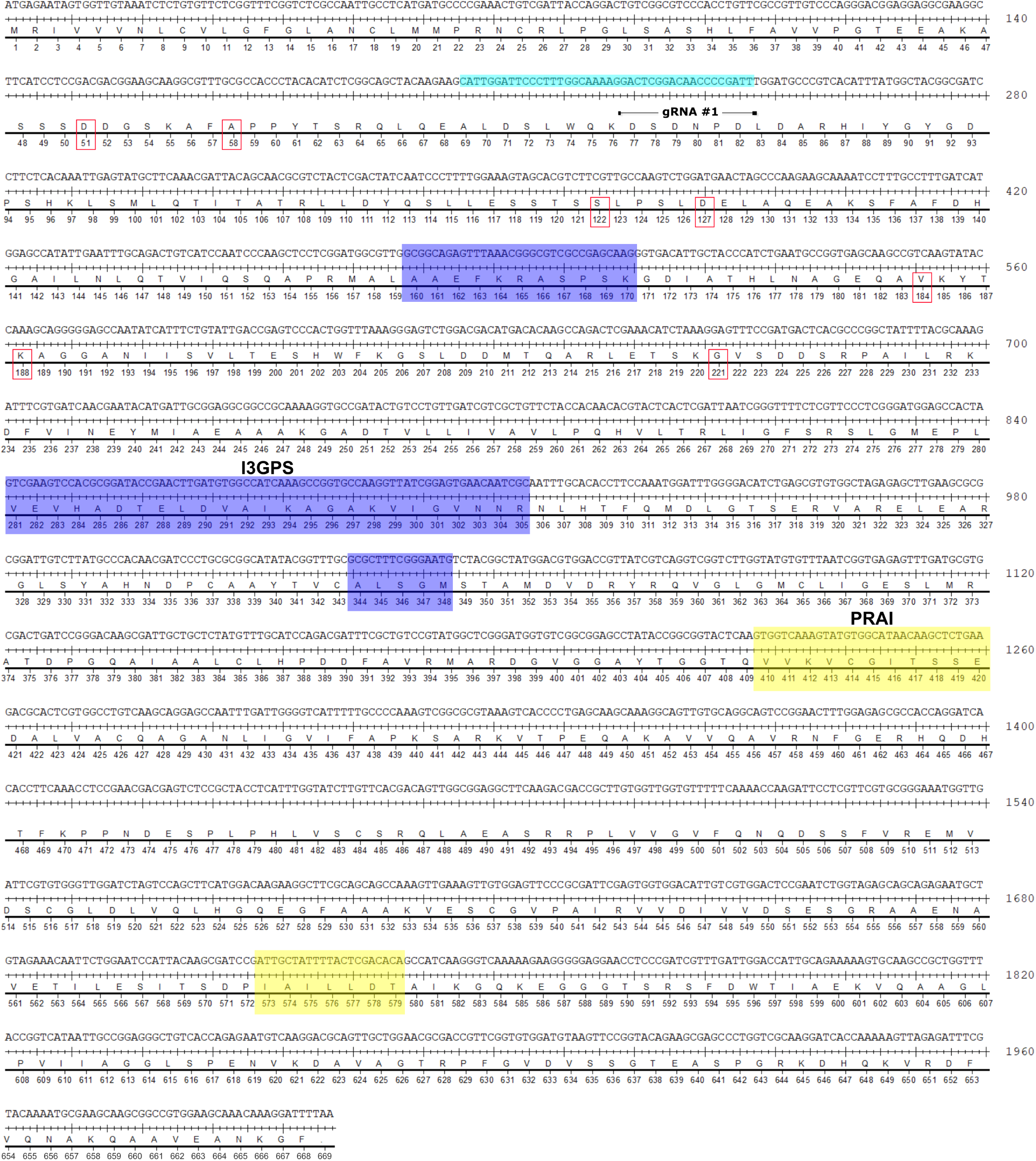
The PtI3GPS-PRAI coding sequence and gRNA target sites. The coding regions for the I3GPS (blue) and PRAI (yellow) catalytic residues are highlighted. Residues that differ between the two alleles are indicated by red boxes.

**Figure 8.**
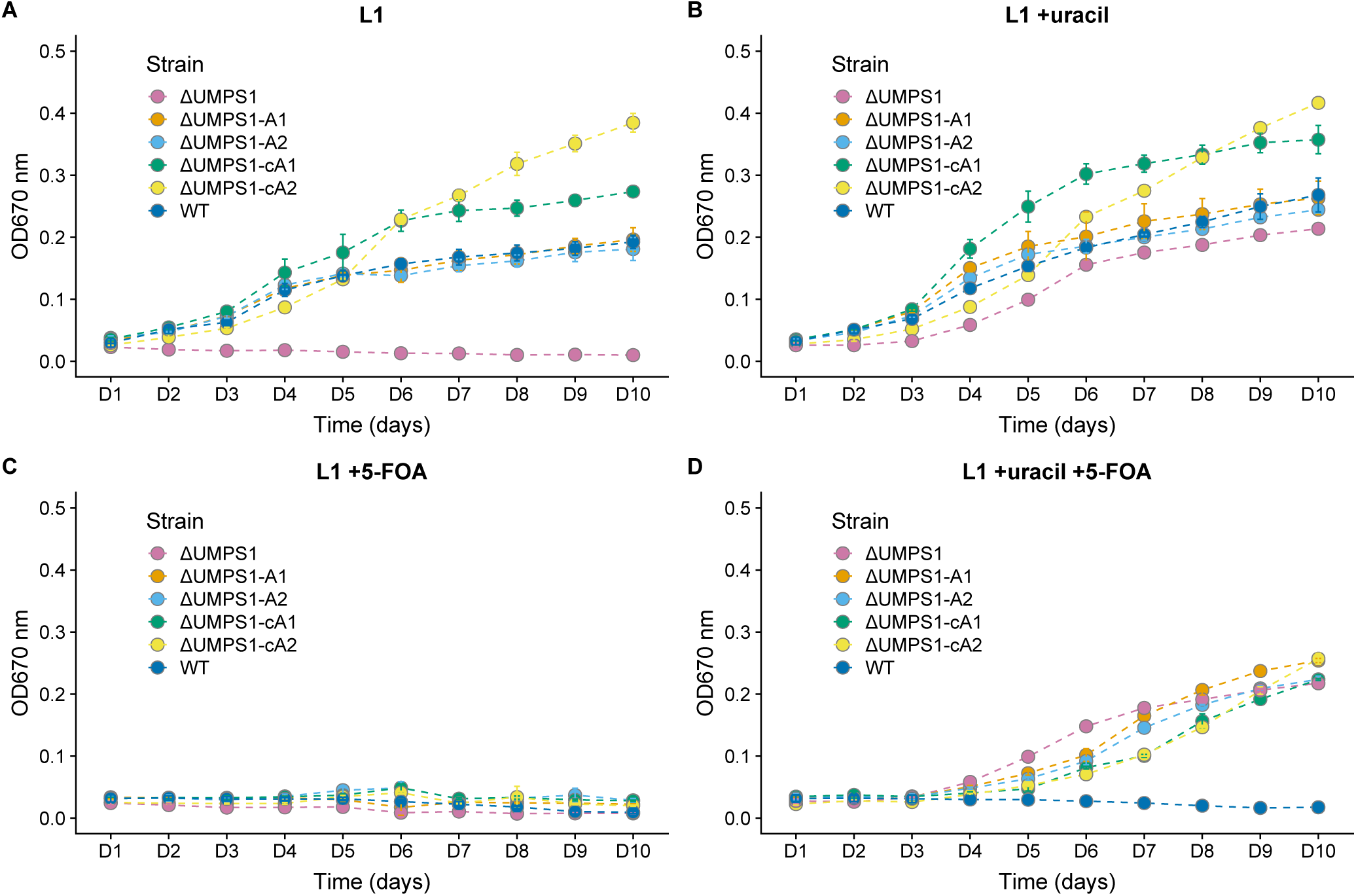
Assaying ΔUMPS1 knockout and complemented growth rates in L1 media supplemented with uracil, 5-FOA, or both. A biallelic PtUMPS mutant (ΔUMPS1) is unable to grow in L1 media without uracil supplementation. Reintroducing the gene on a stably replicating plasmid restores the WT phenotype in L1 media. (A) Growth rates in L1 media. (B) Growth rates in L1 media supplemented with uracil. (C) Growth rates in L1 media supplemented with 5-FOA. (D) Growth rates in L1 media supplemented with uracil and 5-FOA. WT, Wild-type *P. tricornutum*; ΔUMPS1, PtUMPS knockout strain 1; ΔUMPS1-A1, ΔUMPS1 possessing pPtUMPSA1; ΔUMPS1-A2, ΔUMPS1 possessing pPtUMPSA2; ΔUMPS1-cA1, ΔUMPS1 possessing pPtUMPScA1; ΔUMPS1-cA2, ΔUMPS1 possessing pPtUMPScA2. Points represent mean values and error bars represent standard deviation for three replicates.

**Figure 9.**
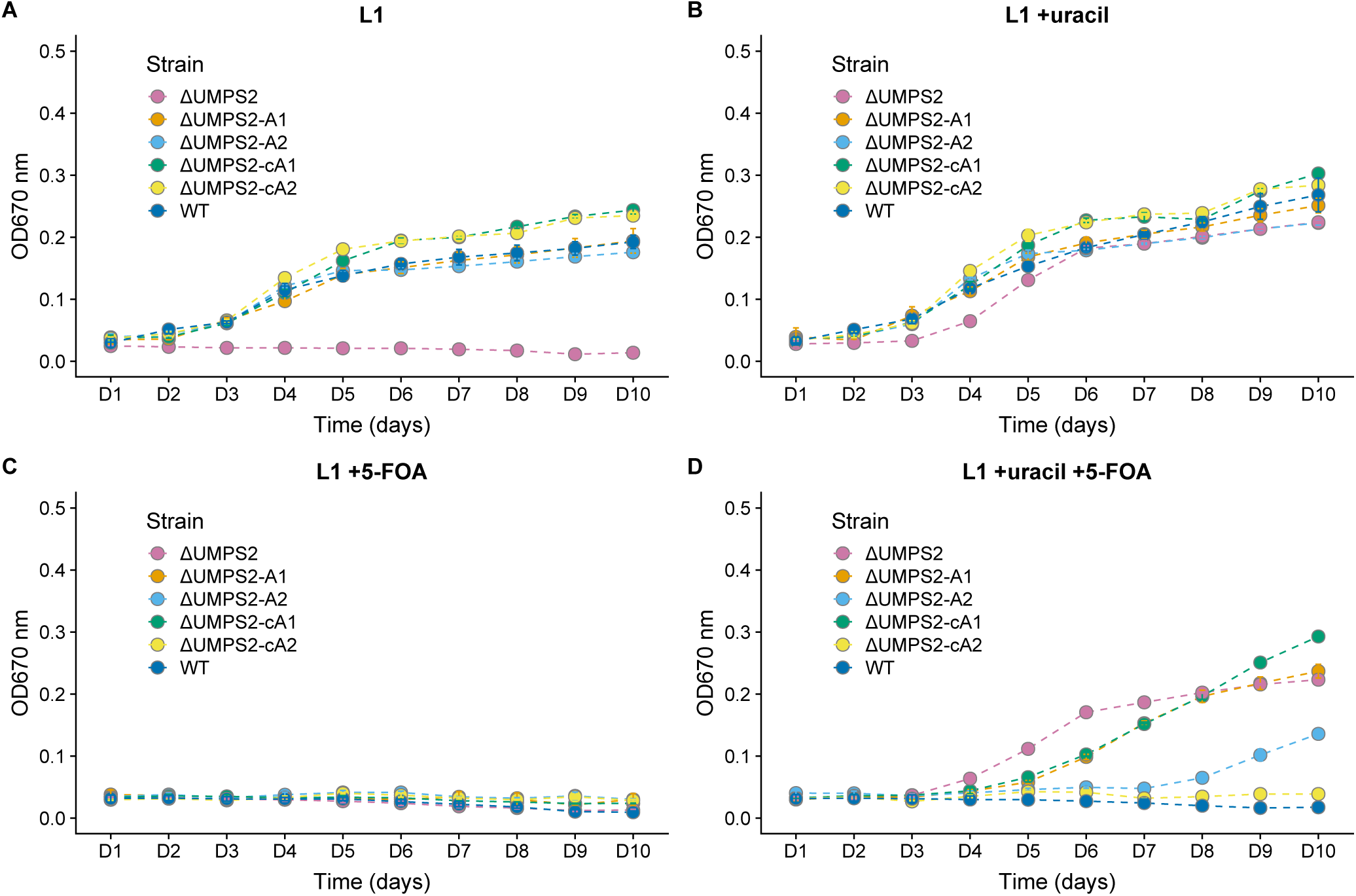
Assaying ΔUMPS2 knockout and complemented growth rates in L1 media supplemented with uracil, 5-FOA, or both. A biallelic PtUMPS mutant (ΔUMPS2) is unable to grow in L1 media without uracil supplementation. Reintroducing the gene on a stably replicating plasmid restores the WT phenotype in L1 media. (A) Growth rates in L1 media. (B) Growth rates in L1 media supplemented with uracil. (C) Growth rates in L1 media supplemented with 5-FOA. (D) Growth rates in L1 media supplemented with uracil and 5-FOA. WT, Wild-type *P. tricornutum*; ΔUMPS2, PtUMPS knockout strain 1; ΔUMPS2-A1, ΔUMPS2 possessing pPtUMPSA1; ΔUMPS2-A2, ΔUMPS2 possessing pPtUMPSA2; ΔUMPS2-cA1, ΔUMPS2 possessing pPtUMPScA1; ΔUMPS2-cA2, ΔUMPS2 possessing pPtUMPScA2. Points represent mean values and error bars represent standard deviation for three replicates.

**Table 1.**
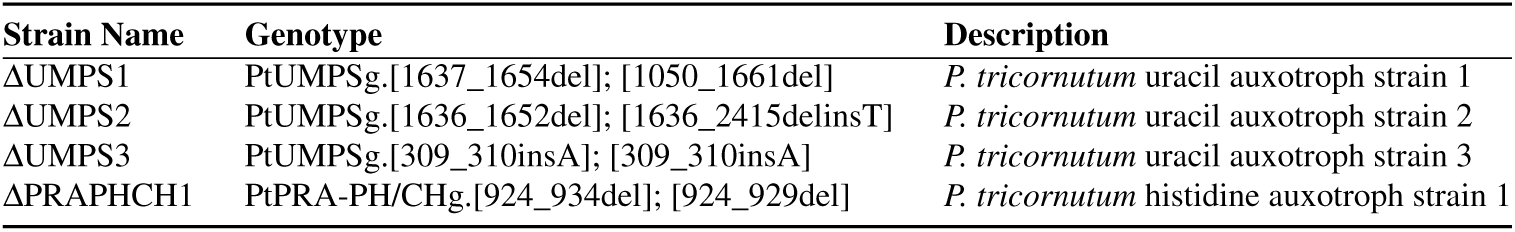
List of *P. tricornutum* auxotroph genotypes.

| Strain Name | Genotype | Description |
| --- | --- | --- |
| ΔUMPS1 | PtUMPSg.[1637_1654del]; [1050_1661del] | *P. tricornutum* uracil auxotroph strain 1 |
| ΔUMPS2 | PtUMPSg.[1636_1652del]; [1636_2415delinsT] | *P. tricornutum* uracil auxotroph strain 2 |
| ΔUMPS3 | PtUMPSg.[309_310insA]; [309_310insA] | *P. tricornutum* uracil auxotroph strain 3 |
| ΔPRAPHCH1 | PtPRA-PH/CHg.[924_934del]; [924_929del] | *P. tricornutum* histidine auxotroph strain 1 |

**Table 2.** List of plasmids used in this study.

| Plasmid | Description | Reference or Source |
| --- | --- | --- |
| pPtGE31 | *P. tricornutum* expression vector | Slattery, et al., 2018 |
| pPtGE34 | *P. tricornutum* expression vector, 40SRPS8 promoter and terminator driving Sh ble, FcpB promoter and FcpA terminator driving Cas9 | Slattery, et al., 2018 |
| pPtGE35 | *P. tricornutum* expression vector, 40SRPS8 promoter and terminator driving Sh ble, FcpB promoter and FcpA terminator driving TevCas9 | Slattery, et al., 2018 |
| pPtUMPSA1 | pPtGE31 encoding PtUMPS allele 1 driven by the PtUMPS promoter and terminator | This study |
| pPtUMPSA2 | pPtGE31 encoding PtUMPS allele 2 driven by the PtUMPS promoter and terminator | This study |
| pPtUMPScA1 | pPtGE31 encoding PtUMPS allele 1 cDNA driven by the PtUMPS promoter and terminator | This study |
| pPtUMPScA2 | pPtGE31 encoding PtUMPS allele 2 cDNA driven by the PtUMPS promoter and terminator | This study |
| pPtPRAPHCH | pPtGE31 encoding PtPRA-PH/CH allele 1 driven by the PtPRA-PH/CH promoter and terminator | This study |
| pTA-Mob | Mobilization helper plasmid required for conjugation | Strand, et al., 2014 |

**Table 3.**
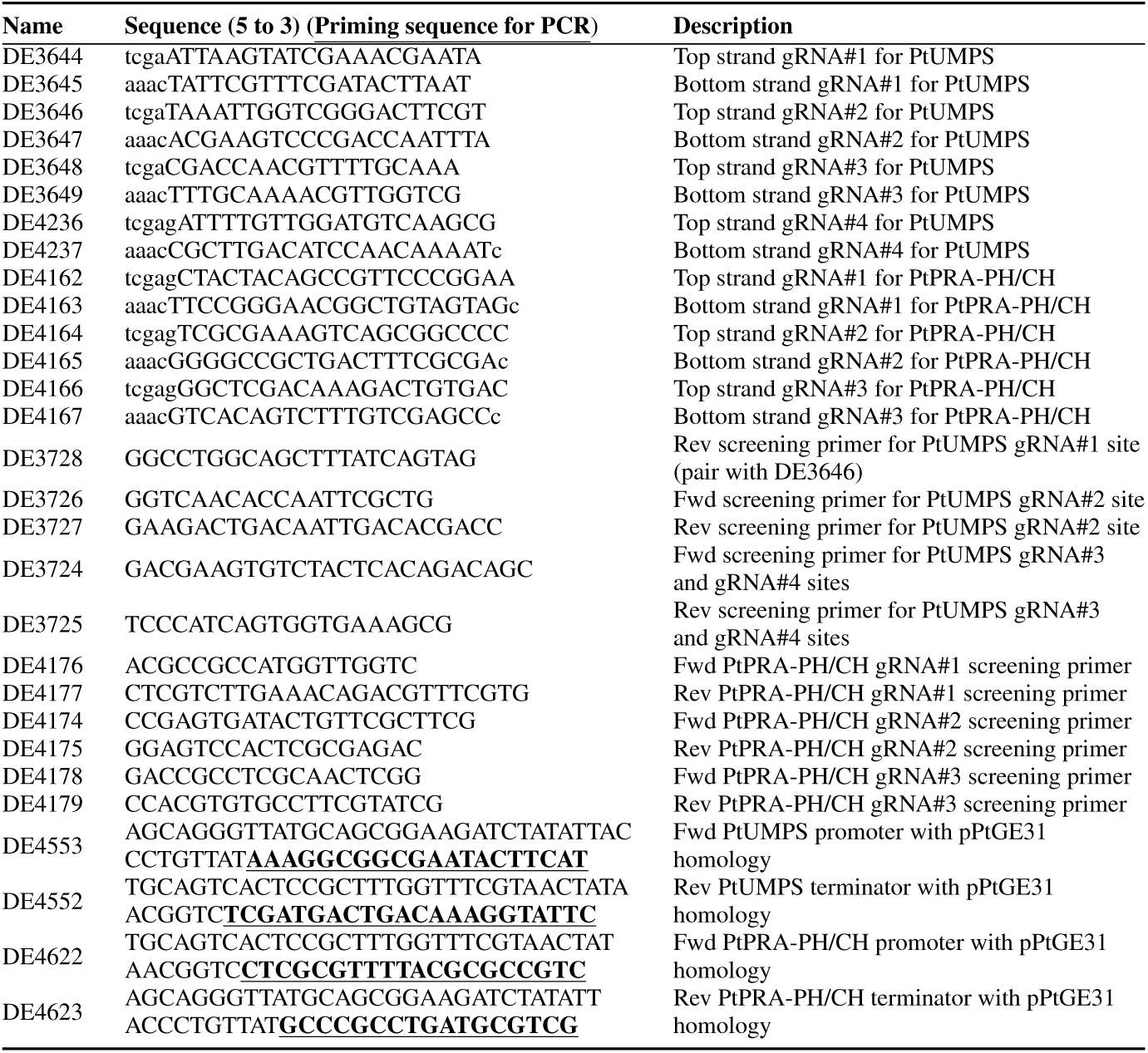
List of oligonucleotides used in this study.

**Table 4.**
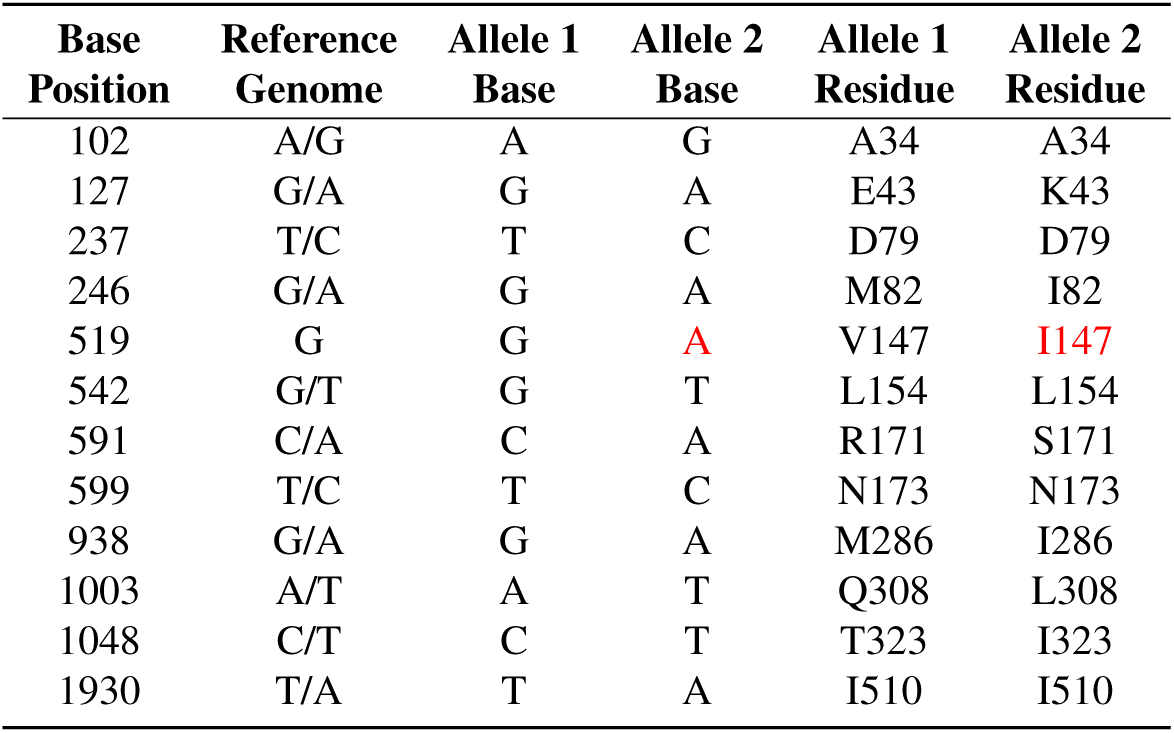
Wild-type *P. tricornutum* UMPS sequence analysis. Base positions are numbered relative to the first base of the start codon in the genomic PtUMPS sequence. SNPs highlighted in red were not present in the reference genome. SNPs located in intronic sequences were not included in this table.

**Table 5.**
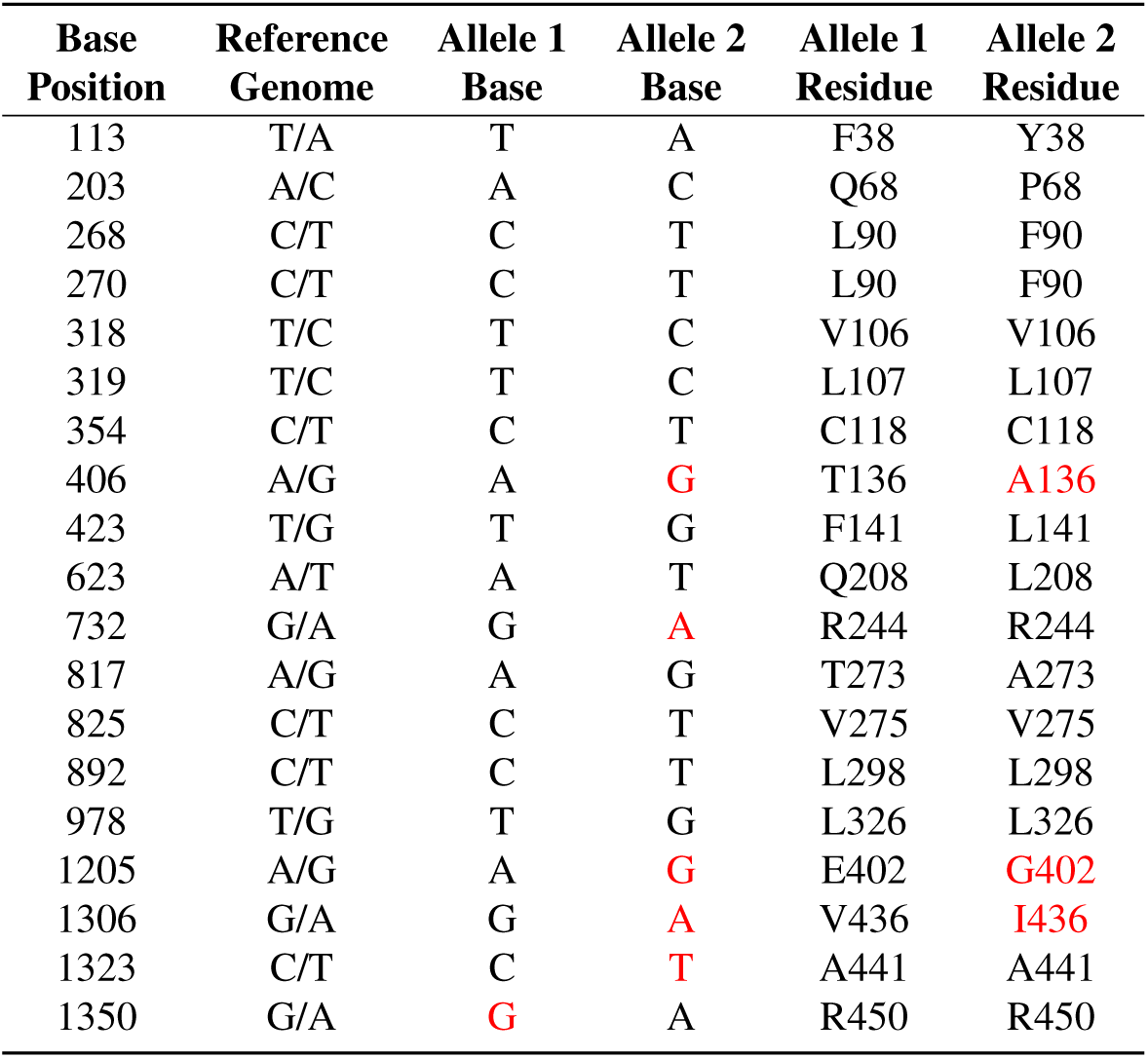
Wild-type *P. tricornutum* PRA-PH/CH sequence analysis. Base positions are relative to the first base of the start codon in the genomic PtPRA-PH/CH sequence. SNPs highlighted in red were not present in the reference genome. Base positions are numbered starting from the first base of the start codon.

**Table 6.** Growth rates of WT and UMPS complement strains in minimal L1 media. Generation times represent the mean value standard deviation for three replicates.

| *P. tricornutum* strain | generation time (hours) |
| --- | --- |
| WT | 29.8 2.6 |
| ΔUMPS2 + pPtUMPSA1 | 27.9 2.8 |
| ΔUMPS2 + pPtUMPSA2 | 24.8 1.5 |
| ΔUMPS2 + pPtUMPScA1 | 28.5 0.9 |
| ΔUMPS2 + pPtUMPScA2 | 23.4 0.8 |
| ΔUMPS1 + pPtUMPSA1 | 26.5 2.7 |
| ΔUMPS1 + pPtUMPSA2 | 29.8 0.7 |
| ΔUMPS1 + pPtUMPScA1 | 26.5 4.0 |
| ΔUMPS1 + pPtUMPScA2 | 30.7 1.2 |

